# Proteomic analysis of necroptotic extracellular vesicles

**DOI:** 10.1101/2020.04.11.037192

**Authors:** Inbar Shlomovitz, Gali Yanovich-Arad, Ziv Erlich, Liat Edry-Botzer, Sefi Zargarian, Hadar Cohen, Yifat Ofir-Birin, Neta Regev-Rudzki, Motti Gerlic

## Abstract

Necroptosis is a regulated and inflammatory form of cell death. We, and others, have previously reported that necroptotic cells release extracellular vesicles (EVs). We have found that necroptotic EVs are loaded with proteins, including the phosphorylated form of the key necroptosis-executing factor, mixed lineage kinase domain-like kinase (MLKL). However, neither the exact protein composition, nor the impact, of necroptotic EVs have been delineated. To characterize their content, EVs from necroptotic and untreated U937 cells were isolated and analyzed by mass spectrometry-based proteomics. A total of 3337 proteins were identified, sharing a high degree of similarity with exosome proteome databases, and clearly distinguishing necroptotic and control EVs. A total of 352 proteins were significantly upregulated in the necroptotic EVs. Among these were MLKL and caspase-8, as validated by immunoblot. Components of the ESCRTIII machinery and inflammatory signaling were also upregulated in the necroptotic EVs, as well as currently unreported components of vesicle formation and transport, and necroptotic signaling pathways. Moreover, we found that necroptotic EVs can be phagocytosed by macrophages to modulate cytokine and chemokine secretion. Finally, we uncovered that necroptotic EVs contain tumor neoantigens, and are enriched with components of antigen processing and presentation. In summary, our study reveals a new layer of regulation during the early stage of necroptosis, mediated by the secretion of specific EVs that influences the microenvironment and may instigate innate and adaptive immune responses. This study sheds light on new potential players in necroptotic signaling and its related EVs, and uncovers the functional tasks accomplished by the cargo of these necroptotic EVs.

## Introduction

Throughout life, cell death is a key element in a vast range of essential biological processes, from embryonic development and organ function to full-body homeostasis and immune responses against tissue injury, pathogens, and tumorigenesis^1^. Necroptosis is a well-studied form of regulated necrosis, defined as receptor-interacting serine/threonine-protein kinase 3 (RIPK3)-/mixed lineage kinase domain-like (MLKL)-dependent, caspase-independent cell death^2–6^. Activation of death receptors, Toll-like receptors (TLRs), or intracellular receptors engages regulated cell death machineries. In these circumstances, caspase-8, together with FLICE-like inhibitory protein (FLIP), cleave and inactivate RIPK1 and RIPK3 to block necroptosis^7–13^.

The ligation of tumor necrosis factor-*α* (TNF-*α*) to TNF receptor 1 (TNFR1), for example, recruits TNFR1-associated death domain (TRADD) and RIPK1 to assemble Complex I, together with TNF receptor associated factor 2 (TRAF2), cellular inhibitor of apoptosis proteins (cIAPs), and linear ubiquitin chain assembly complex (LUBAC)^14–18^. In this complex, RIPK1 ubiquitination drives nuclear factor kappa-light-chain enhancer of activated B cells (NF-κB) signaling resulting in pro-inflammatory and pro-survival signals^19–21^. When Complex I is impaired, *i.e*., via inhibition of cIAPs by second mitochondria-derived activator of caspases (SMAC) mimetics, Complex II is assembled, containing RIPK1, caspase-8, FLIP, and FAS-associated death domain protein (FADD). This can result in caspase-8 cleavage and activation and, finally, apoptosis^17,22–24^.

However, caspase-8 suppression by pharmacologic agents, pathogen effectors, or genetical manipulation unleashes RIPK1 and RIPK3 inhibition. Consequently, auto- and trans-phosphorylation between RIPK1 and RIPK3 leads to the aggregation and phosphorylation of MLKL by RIPK3^8,25–27^. Phosphorylated MLKL (pMLKL) then translocates to the plasma membrane to compromise membrane integrity and execute necroptosis^28–30^. Necroptosis is morphologically similar to necrosis, featuring cell swelling and membrane permeabilization, and driving the release of danger associated molecular patterns (DAMPs), such as interleukin-33, ATP, high mobility group box 1 (HMGB1), and more^2,5,31–33^. Hence, necroptosis is considered an inflammatory form of cell death.

As such, necroptosis has been suggested to contribute to different inflammatory pathologies, including ischemia-reperfusion renal injury^34^, pancreatitis^8^, atherosclerosis^35^, and neurodegenerative diseases^36–39^. In addition, necroptosis is reported to be involved in host-pathogen interactions. Herpes simplex virus (HSV) induces necroptosis in mice, but prevents necroptosis in humans, which are its natural host^40–42^. Many other viruses target and block necroptosis, such as vaccinia virus, cytomegalovirus (CMV), Epstein-Barr virus (EBV), and Influenza A^43–48^. In contrast, few bacteria have been shown to induce necroptosis. Among those that do are *Salmonella enterica, Mycobacterium tuberculosis*, and *Staphylococcus aureus*^49–53^. The inflammation-inducing quality of necroptosis has recently been applied to cancer research. Vaccination with necroptotic cancer cells was shown to induce anti-tumor immunity by releasing DAMPs and inducing maturation of antigen-presenting cells (APCs) and tumor-antigen loading^54–58^. Cross-priming of cytotoxic T cells by necroptotic cancer cells has also been reported to require their RIPK1-mediated NF-κB signaling^59^.

In contrast to necroptosis, apoptosis is classified as an immunologically silent form of cell death, characterized by plasma membrane blebbing^60^ and the shedding of apoptotic bodies^61–63^. A key feature of apoptosis is the exposure of phosphatidylserine (PS) on the outer plasma membrane, which functions as an “eat me” signal, resulting in phagocytosis and clearance of apoptotic cells and bodies. We^64^ and others^65,66^ have previously revealed that necroptotic cells also expose PS to the outer plasma membrane prior to its rupture. We discovered that PS-exposing necroptotic cells release extracellular vesicles (EVs), which are smaller and contain a higher protein content than apoptotic bodies^64^. Endosomal sorting complexes required for transport (ESCRT) machinery is a family of proteins that plays a role in endosomal protein transport, multivesicular endosome (MVE) formation, and budding^67^. By using a dimerizable RIPK3 or MLKL system, Gong *et al*. revealed ESCRTIII-dependent shedding of PS-exposing plasma membrane bubbles during necroptosis. ESCRTIII-mediated membrane shedding during necroptosis delayed cell death and enabled longer inflammatory signaling^65,68^. These results underline a major knowledge gap, as the exact protein content, as well as the biogenesis and impact, of necroptotic EVs, have not yet been elucidated.

EVs are cell-derived membranous structures, divided into exosomes, which are 50-150 nm in size, and microvesicles, which are 50-500 nm^69^. Exosomes are generated via the endosomal system as intraluminal vesicles (ILVs) that are formed during the maturation of an MVE and secreted by MVE fusion with the plasma membrane^70,71^. Microvesicles originate from the direct shedding of microdomains of the plasma membrane^72,73^. Yet, in recent years it has become clear that EVs are a heterogeneous group of multiple subgroups, which differ in size and cargo components^74^. While originally thought to function mainly in the elimination of cellular waste, it is now understood that the fate of EVs is much more complex, mediating cell-to-cell communication in numerous settings and mechanisms^69,70,75^.

Overall, this highlights the need for a large-scale proteomic analysis of necroptotic EVs to define both the upstream necroptotic mechanisms and the downstream effects of these EVs. To this end, we utilized mass spectrometry-based proteomics to characterize EVs extracted from necroptotic U937 cells compared to control EVs from untreated cells. This study sheds light on new potential players in necroptotic signaling and suggests a new mechanism mediating necroptosis-induced inflammation.

## Materials and Methods

### Cell culture

The U937 human histiocytic lymphoma cell-line^76^ was cultured in RPMI-1640 (Biological industries), supplemented with 10 % fetal bovine serum (FBS) (Gibco), 1 % penicillin-streptomycin (Gibco) and 10mM HEPES (Gibco), at 37 °C in a humidified 5 % CO_2_ atmosphere.

### Cell death stimuli

U937 cells were seeded at 1 × 10^6^ cells/mL in FBS-free, EVs-depleted, RPMI medium and were treated with TNF-*α*, Birinapant, and QVD-OPh (denoted TBQ) or with TNF-*α*, second mitochondria-derived activator of caspases (SMAC mimetic, AZD5582), and z-VAD-fmk (denoted TSZ) to induce necroptosis (Table 1), or left untreated. When indicated, cells were treated with TNF-*α* alone (denoted T) as a control, or with TNF-*α* and Birinapant (denoted TB) to induce apoptosis.

**Table 1.** Concentrations of reagents used to induce cell death.

| Reagent | Concentration | Manufacture |
| --- | --- | --- |
| TNF-α | 1.15 nM | PeproTech |
| AZD5582 | 2.5 μM | Tocris Bioscience |
| Birinapant | 2 μM | APExBIO |
| z-VAD-fmk | 20 μM | Calbiochem |
| QVD-OPh | 20 μM | APExBIO |

### Cell staining

When indicated, additional staining with propidium iodide (PI) and AnnexinV (A5) was performed (Table 2). A5 was added directly to the medium for live-cell imaging, or was added in A5 binding buffer (eBioscience) for 10 min at room temperature before washing for flow cytometry analysis. For carboxyfluorescein succinimidyl ester (CFSE) staining, prior to addition of cell death stimuli, cells were washed twice in PBS, resuspended at 5 × 10^6^ cells/mL in PBS containing CFSE, and incubated for 10 min at 37 °C. Subsequently, five volumes of ice-cold medium were added, and cells were incubated for 5 min at 4 °C, followed by three washes, and a final resuspension in medium. Hoescht33342 was added directly to the medium.

**Table 2.** Target, dilutions and concentrations of reagents used for staining.

| Reagent | Target | Permeability | Dilution/Concentration | Manufacture |
| --- | --- | --- | --- | --- |
| Propidium Iodide (PI) | Nucleic acids | - | 1 μg/mL | Sigma-Aldrich |
| AnnexinV (A5) | Phosphatidylserine | - | 1:200 | MBL |
| Carboxyfluorescein succinimidyl ester (CFSE) | Cellular amines (lysine residues) | + | 5 μM | Sigma-Aldrich |
| Hoescht33342 | Nucleic acids | + | 1 μg/mL | Sigma-Aldrich |

### Cell viability assessment

To assess cell viability, 100 μL of treated and untreated cells, supplemented with PI and A5, as described above, were plated in 96-well plates in triplicate. For live-cell imaging, plates were placed on the IncucyteZOOM (Essen BioScience) and were recorded every 10-30 min. Data were analyzed using the IncucyteZoom2016B analysis software and exported to GraphPad Prism software.

Supplementary assessment was performed using flow cytometry. Treated and untreated cells were collected into a 96-well U-shaped plate. Cells were stained with PI and A5 as described above and re-suspended in 200 μL PBS. Samples were acquired by the flow cytometry using the Attune NxT (Thermo Fisher Scientific) and data were analyzed using Flowjo software (TreeStar).

### Extracellular vesicle extraction

2 × 10^7^ U937 cells per group were stimulated and assessed for cell death, as above. When TBQ-treated U937 cells reached 60 % PI-positivity by live-cell imaging and flow cytometry (4 h post TSZ-stimuli or 5–6.5 h post TBQ-stimuli), supernatants were collected and centrifuged at 400 × g for 5 min to pellet cells. Supernatants were further centrifuged at 1500 × g for 10 min to remove debris. Supernatants were then centrifuged at 14,000 × g for 70 min to remove micro-vesicles and filtered through a 0.45 μm filter. Finally, supernatants were centrifuged at 100,000 × g for 190 min in ultracentrifugation using type 70 Ti rotor (Oprima XL-80K, Beckman) to pellet the EVs (Fig. 1E). For proteomic analysis, six pairs of independently extracted EVs from untreated and TBQ-treated cells were analyzed as biological replicates by mass spectrometry. For validation by immunoblot analysis, EVs were extracted in two additional independent experiments.

**Figure 1.**
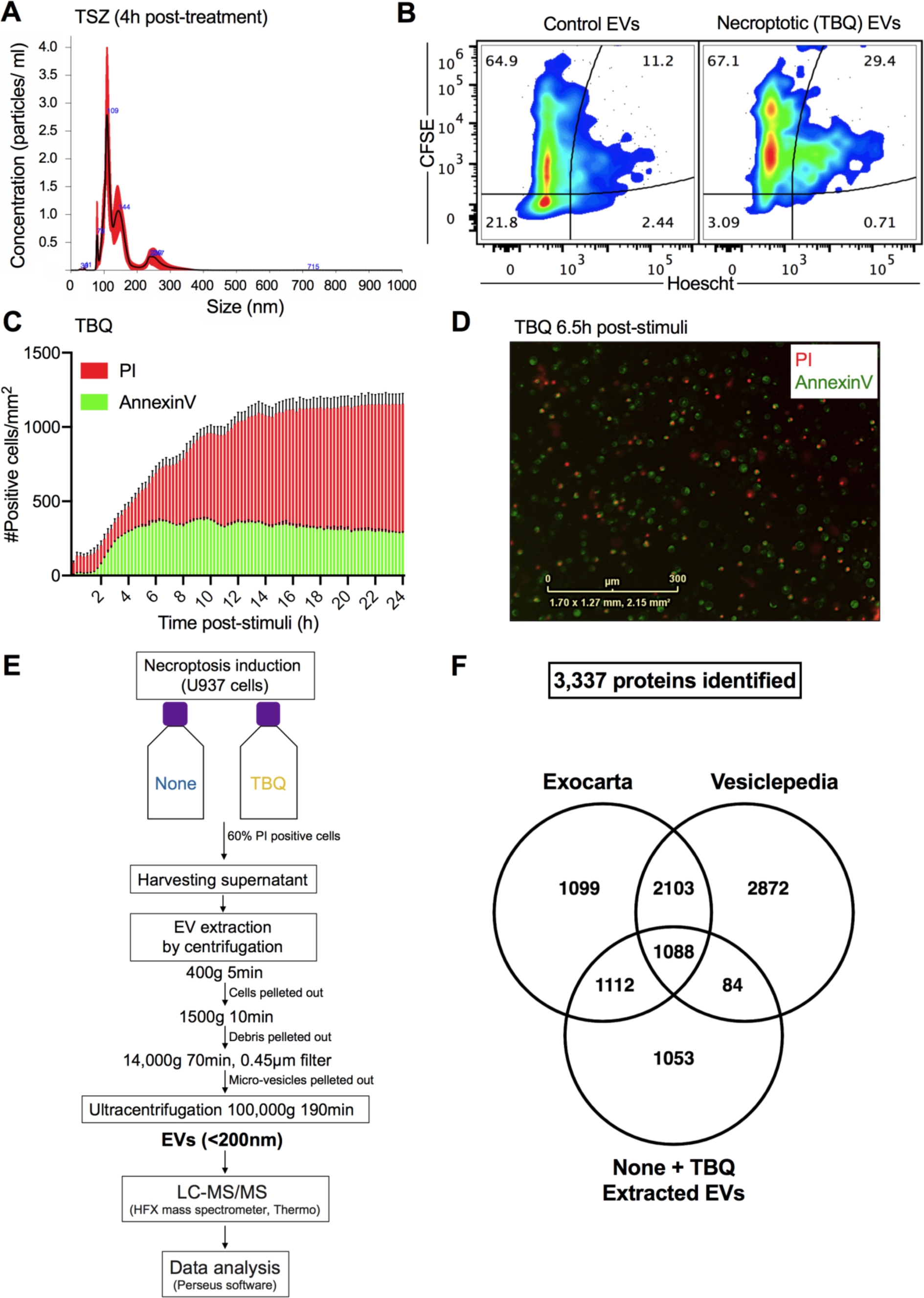
Extraction of necroptotic extracellular vesicles (EVs) **A**, U937 cells were treated with TNF-*α* (1.15 nM), SMAC (AZD5582, 2.5 mM), and z-VAD-fmk (20 mM) (denoted TSZ) to induce necroptosis. After 4 h, EVs were extracted using ultracentrifugation and analyzed for size and concentration by NanoSight. **B**, CFSE-stained U937 cells were treated with TNF-*α* (1.15 nM), Birinapant (5 mM), and QVD-OPh (20 mM) (denoted TBQ) to induce necroptosis. Five hours later, EVs were extracted using ultracentrifugation, stained with Hoescht33342, and analyzed by flow cytometry (Attune NxT) for CFSE and Hoescht fluorescence intensity to examine their protein and nucleic acid composition, respectively. **C, D**, U937 cells were treated with TBQ as above to induce necroptosis. Cell viability was monitored using AnnexinV-FITC (green) and PI (red) via real-time imaging (IncucyteZoom). **C**, Number of AnnexinV- and PI-positive cells per mm2 at the indicated time-points post-stimuli. **D**, Representative image of treated cells 6.5 h post-stimuli. **E**, Schematic overview of the experimental and data analysis procedure. U937 cells were left untreated as a control (denoted none) or stimulated for necroptosis (TBQ). When TBQ-treated cells reached 60 % PI-positivity, supernatants were harvested for serial centrifugation and ultracentrifugation. Six pairs of independently extracted EVs were analyzed as biological replicates by mass spectrometry and Perseus data analysis software. **F**, 3,337 proteins were identified in the extracted EVs from both control (none) and necroptotic (TBQ-treated) cells by mass spectrometry. Venn diagram of total proteins identified in either none- or TBQ-extracted EVs compared with the exosome proteome data bases, Exocarta and Vesiclepedia. **A, C, D**, Data are representative of at least three independent experiments. **A**, Data are presented as the mean of five acquired samples ± SEM and representative of three independent experiments. **B**, Plots are representative of duplicate samples. **C**, Data are presented as the mean of triplicate samples ± SD. EVs, extracellular vesicles; TSZ, TNF-*α*, SMAC and z-VAD-fmk; TBQ, TNF-*α*, Birinapant and QVD-OPh; PI, propidium iodide; h, hours; LC-MS/MS, liquid chromatography with tandem mass spectrometry; SEM, standard error of the mean; SD, standard deviation.

### NanoSight EV analysis

For a single particle concentration and size measurements, EVs were extracted from TSZ-treated U937 cells four h post-stimuli and analyzed using Nanoparticle Tracing Analysis (NTA) (Malvern Instruments, NanoSight NS300, version 3.1.54).

### In gel proteolysis and mass spectrometry analysis

Pelleted EVs were suspended in 50 μL sodium dodecyl sulphate-polyacrylamide gel electrophoresis (SDS-PAGE) gel loading buffer (buffer composition is described below) and samples were heated at 95 °C for 5 min immediately before loading. Samples were loaded into SDS-PAGE precast gels (BIO-RAD). The proteins in the gel were reduced with 2.8mM DTT (60 °C for 30 min), modified with 8.8 mM iodoacetamide in 100 mM ammonium bicarbonate (in the dark, room temperature for 30 min) and digested in 10 % acetonitrile and 10mM ammonium bicarbonate with modified trypsin (Promega) at a 1:10 enzyme-to-substrate ratio, overnight at 37 _°_C. An additional second trypsinization was done for 4 h. The resulting tryptic peptides were resolved by reverse-phase chromatography on 0.075 × 200-mm fused silica capillaries (J&W) packed with Reprosil reversed phase material (Dr Maisch GmbH, Germany). The peptides were eluted with linear 105 min gradient of 5 % to 28 % acetonitrile with 0.1 % formic acid in water, 15 min gradient of 28 % to 90 % acetonitrile with 0.1 % formic acid in water, and 15 min at 90 % acetonitrile with 0.1 % formic acid in water at flow rates of 0.15 μl/min. Mass spectrometry was performed by a Q-Exactive plus mass spectrometer (QE, Thermo) in a positive mode using repetitively full MS scan followed by High energy Collision Dissociation (HCD) of the 10 most dominant ions selected from the first MS scan.

The mass spectrometry data from the biological repeats were analyzed using the MaxQuant software 1.52.8 vs. the human part of the Uniprot database with 1 % FDR in the peptide-spectrum match (PSM) and protein level^77,78^. The data were quantified by maxLFQ algorithm using the same software^79^.

Of note, Coomassie blue staining demonstrated similar stain of the control and the necroptotic EV samples, suggesting similar protein concentration among both groups (supplementary Fig. S1). This is supported by the fact that LFQ intensity normalization was successfully performed.

### Proteomic data analysis

Statistical analysis of the identification and quantization results was done on 3,384 protein groups using Perseus software^80^, filtered into 3,337 unique gene names.

Venn diagrams were generated using Venny 2.1.0 tool^81^ (https://bioinfogp.cnb.csic.es/tools/venny/index.html) against the exosome proteome data bases, Exocarta^82–84^ (http://www.exocarta.org/) and Vesiclepedia^85,86^ (http://microvesicles.org/index.html). Each data set was selected for the human proteins, followed by filtering of duplicates.

Total proteins identified in necroptotic EVs were defined as proteins with valid quantitative values in at least three replicates of the TBQ samples.

In all analyses, values were log_2_-transformed and missing values imputation per sample was performed by drawing values from a normal distribution with a width of 0.5 of the sample’s valid standard deviation and a downshift of 1.4 standard deviations. LFQ intensity mentioned in text and figures are in log_2_ scale. Unless otherwise mentioned, fold change represents the raw ratio between the mean LFQ intensity values in necroptotic and control EVs.

For the unsupervised clustering analyses and principal component analysis, a minor batch effect due to sample preparation time points was removed using the R limma package^87^. For the hierarchical clustering based on protein expression, data were filtered to retain only proteins with valid quantitative values in three replicates of at least one group, resulting in 2,984 proteins in total.

For the supervised analysis, one side Student’s t-test for paired samples was used with a false discovery rate (FDR) cut-off of 0.1 and S0=0.1 to define “TBQ-upregulated proteins” to be used as the significantly upregulated proteins in necroptotic EVs in all downstream analysis. For enrichment analysis, a list of EV proteins was downloaded from Vescilepedia database^85,86^ to be used as background against which the t-test significant proteins were tested (Fisher’s exact test, Benjamini-Hochberg FDR < 0.02). To identify protein-protein interactions in the TBQ-upregulated proteins, we analyzed the list of “TBQ-upregulated proteins” using String protein-protein interaction database (https://string-db.org/) and Cytoscape software^88^.

### Data availability

The mass spectrometry proteomics data of 12 samples have been deposited to the ProteomeXchange Consortium^89^ via the PRIDE^90,91^ partner repository with the dataset identifier PXD018258.

### EV lysates for Western blotting

EVs from 2 × 10^7^ treated or untreated U937 cells were extracted as above. Following ultracentrifugation, the supernatants was discarded and the pellet (containing the EVs) was lysed in 40 μL lysis buffer (150 mM NaCl, 1 % (v/v) Triton X-100, 1 % (w/v) sodium deoxycholate, 0.1 % (w/v) SDS in 10 mM Tris-HCl, pH 7.5), supplemented with Halt protease and the phosphatase inhibitor cocktail ethylenediamine tetra-acetic acid (EDTA) free (1:100, Pierce Biotechnology) immediately prior to use. Following 15 min of incubation at 4 °C, lysates were centrifuged at 13000 × g for 20 min at 4 °C. 5× SDS-PAGE gel loading buffer (0.5 M DTT, 10 % (w/v) SDS, 50 % (v/v) glycerol, 0.2 % (w/v) bromophenol blue powder, 10 % (v/v) water in 1 M Tris, pH 6.8) was added to the suspension and samples were heated at 95 °C for 5 min immediately before loading.

### Western blotting

Samples were loaded into SDS-PAGE precast gels (BIO-RAD) for Western blot analysis. Proteins were transferred onto 0.2 μm nitrocellulose membrane using the Trans-Blot Turbo Transfer system (BIO-RAD). Membranes were blocked with 5 % skim milk in TBS-T (20 mM Tris pH 7.4, 150 mM NaCl, 0.05 % Tween-20) for 1 h and probed overnight with the primary antibodies (Table 3) diluted in 5 % skim milk in TBS-T. The next day, membranes were washed five times before the horseradish peroxidase (HRP)-conjugated secondary antibodies (Table 4) (diluted as above) were added for 1 h. Images were taken using Odyssey Fc system (LI-COR Biosciences) and analyzed using ImageStudio analysis software (LI-COR Biosciences).

**Table 3.**
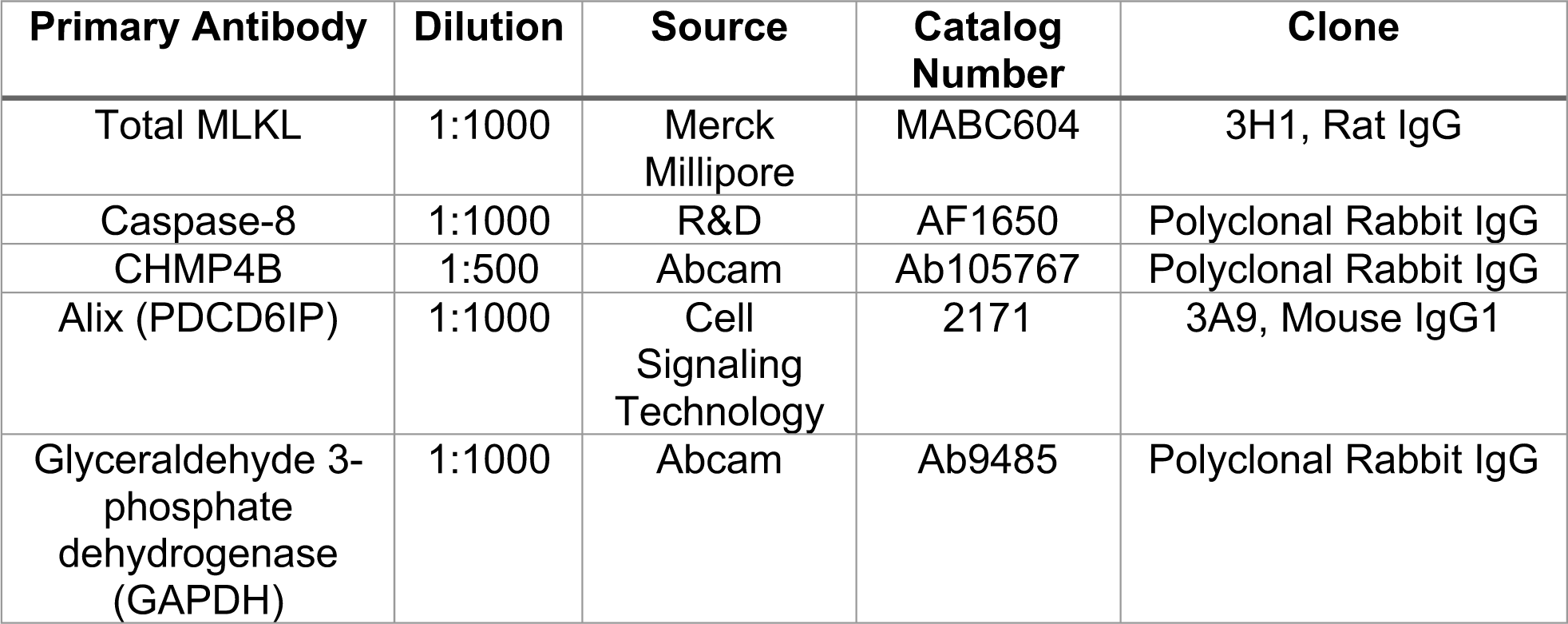
Primary antibodies used for Western blots.

**Table 4.** Secondary antibodies used for Western blots.

| Secondary Antibody | Dilution | Source | Catalog Number |
| --- | --- | --- | --- |
| HRP conjugated goat anti-rat | 1:5000 | Jackson Immuno Research Laboratories | 112-035-003 |
| HRP conjugated goat anti-rabbit | 1:5000 | Jackson Immuno Research Laboratories | 111-035-003 |
| HRP conjugated donkey anti-mouse | 1:5000 | Jackson Immuno Research Laboratories | 715-035-150 |

### Mice

C57BL/6J-RccHsd mice were obtained from Harlan Laboratories and grown in-house. All experiments were reviewed and approved by the Animal Care Committee of Tel Aviv University (Number 01-16-105) and were performed according to their regulations and guidelines regarding the care and use of animals for experimental procedures. Experiments were conducted in pathogen-free facilities at Tel Aviv University.

### Purification of thioglycolate-derived peritoneal macrophages

Mice were injected intraperitoneally with 3 mL of 3 % (v/v) thioglycolate (thioglycolic acid, Sigma-Aldrich, T3758) in PBS to elicit peritoneal macrophages using a 25G needle. Three days later, the mice were humanely euthanized and 5 mL of PBS was injected into the peritoneal cavity of each mouse with a 21G needle and the abdomen was gently massaged for 5-10 seconds. The peritoneal fluid was then collected and centrifuged at 400 × g for 5 min to pellet cells. Cells were seeded in DMEM medium, supplemented with 10 % FBS (Gibco), 1 % penicillin-streptomycin (Gibco) and 10 mM HEPES (Gibco) at a concentration of 1 × 10^6^ cells/ mL, and then seeded in 100 μL per well in 96-wells plate the day before use. Cells were purified and seeded from five different mice, as biological replicates, in two independent experiments.

### Phagocytosis *in vitro* assay

For *in vitro* phagocytosis assays, the plate seeded with peritoneal macrophages was washed with PBS three times to discard non-adherent cells. EVs were extracted from U937 cells previously treated with either TNF-*α* alone (T), TNF-*α* and Birinapant (TB), TNF-*α*, Birinapant and QVD-OPh (TBQ), or left untreated. 100 μL of EVs extracted from 7.5 × 10^6^ cells were added per well of a biological triplicate of peritoneal macrophages. Following an overnight incubation, the plates were centrifuged, and supernatants were collected to measure the secretion of cytokines and chemokines by ELISA. When indicated, cells were first stained with CFSE prior to addition of cell death stimuli and EV extraction. For assessment of phagocytosis, peritoneal macrophages were harvested, following an overnight incubation, using 3 mL of 3 mM EDTA in PBS added for 10 min at room temperature, and then analyzed by flow cytometry for their CFSE-positivity, demonstrating the uptake of CFSE-stained EVs.

### Enzyme-linked immunosorbent assay (ELISA)

Murine TNF-*α*, murine CCL2, murine IL-10, and murine IL-6 were measured using commercial ELISA kits obtained from Invitrogen, Thermo Fisher Scientific (Lower detection limits: 8, 15, 32 and 4 pg/mL, respectively), according to manufacturer’s protocol.

### Statistical analysis

Any additional statistical analysis was performed using GraphPad Prism 8 software or Python and is detailed for each experiment separately.

## Results

### Necroptotic Phosphatidylserine (PS)-positive EVs share similarity with exosome databases

PS exposure on the outer cell membrane was long thought of as a phenomenon restricted to early apoptosis^92^. We, and others, have challenged this dogma by demonstrating that PS is also exposed on the outer cell membrane during early necroptosis^64–66,93,94^. Similar to apoptotic cells that form PS-exposing apoptotic bodies, we found that necroptotic cells release PS-exposing EVs. These necroptotic EVs are 70-300 nm in size, with a mean of 138.7 ± 14.4 nm, according to nanoparticle tracking analysis (NTA) by the NanoSight (Fig. 1A), corresponding with the size of exosomes (50-150 nm) and microvesicles (50-500 nm)^69^. In addition, protein labeling by CFSE staining and nucleotide labeling by Hoescht staining, revealed that necroptotic EVs are rich with protein cargo that has not yet been thoroughly studied (Fig. 1B). Thus, we aimed to characterize the proteome of necroptotic EVs in a large-scale analysis by utilizing mass spectrometry (MS)-based proteomics.

In order to correctly time the extraction of necroptotic EVs, we first calibrated necroptotic cell death kinetics in U937 cells, a human histiocytic lymphoma cell-line. Cells were treated with TNF-*α*, Birinapant (a SMAC mimetic that inhibits cIAPs), and the pan-caspase inhibitor QVD-OPh (denoted TBQ) to induce necroptosis. To monitor death kinetics, cells were stained with AnnexinV, a cell impermeable dye that binds to PS, and propidium iodide (PI) that binds nucleic acids upon loss of plasma membrane integrity. Consistent with our previously published data, necroptotic U937 cells stain with AnnexinV in a time-dependent manner before becoming PI-positive, confirming that, during necroptosis, PS exposure on the outer plasma membrane precedes membrane rupture (Fig. 1C). In support of this, AnnexinV-single positive cells were detected for 6.5 h following necroptosis induction (Fig. 1D). Based on these data, we chose to schedule the EV extraction for when 60 % of the cells undergoing necroptosis are PI-positive (5–6.5 h post-stimuli), a time-point at which AnnexinV-positivity has plateaued and membrane rupture, *i.e*., PI positivity, is starting to peak.

To investigate the protein composition of necroptotic EVs, U937 cells were either left untreated as a control, or given a necroptotic stimulus, as described above. At the correct time point (according to 60 % PI positivity), supernatants were harvested, and a series of centrifugations were performed to pellet the cells, cell debris, and micro-vesicles. Finally, ultracentrifugation was conducted to pellet the EVs for consequent analysis by liquid chromatography with tandem mass spectrometry (LC-MS/MS). Data were then quantified by a label free quantification (LFQ) approach and analyzed using Perseus software (Fig. 1E). For the complete proteome analysis, six pairs of independently extracted EVs from untreated and TBQ-treated cells were analyzed as biological replicates. Considering the methods used in our previous publication^64^ and this currently used approach, necroptotic EVs were overall purified using either a density-based separation, a size-exclusion method or ultracentrifugation. Purified EVs were then characterized qualitatively and quantitively by NTA, electron microscopy, and flow-cytometry, and for their protein content by western blot and, finally, mass spectrometry. Hence, our results comply with the EV-TRACK consortium requirements of the information necessary to interpret and reproduce EV experiments^95,96^.

A total of 3,337 proteins were identified in the extracted EVs from both control and necroptotic (TBQ) U937 cells. To test our experimental method in light of current published data regarding vesicle and exosome content, we compared the total proteins identified in either control or necroptotic EVs with the human exosome proteome databases, Exocarta^82–84^ and Vesiclepedia^85,86^, using Venn analysis. Out of 3,337 proteins, 2,200 were present in the Exocarta data base, which constitutes 66 % of the total protein content of the extracted EVs. In addition, the extracted EVs shared 1,172 common proteins with the Vesiclepedia database, *i.e*., 35 % of their total protein content (Fig. 1F). This includes 65 of the 75 most frequently identified proteins in both Exocarta and Vesiclepedia (supplementary Fig. S2 and Table S1). Among these are EV biogenesis factors, *e.g*., Clathrin heavy chain 1 (CLTC), the ESCRT accessory proteins programmed cell death 6-interacting protein (PDCD6IP, also known as Alix), and tumor susceptibility gene 101 (TSG101), as well as tetraspanins, such as CD63 and Flotillin-1 (FLOT1), and intracellular EV trafficking proteins from the annexin and Ras-related protein families^69^. Notably, 1053 proteins are unique to our system.

These results support the validity of our EV extraction system, making our data comparable with established vesicle proteome research, and presenting additional unique content to be studied.

### Necroptotic EVs feature a unique proteome signature

EVs are thought to have a role in cell-cell communication and in cell maintenance via the dumping of unwanted cellular content^97,98^. In support, interfering with necroptotic EV release by silencing the ESCRTIII family member, charged multivesicular body protein 2A (CHMP2A), or Rab27a and Rab27b that are required for exocytosis, increases the sensitivity of cells to necroptosis^65,66^. In addition, EVs are known to function in cell-to-cell communication by modifying their recipient cells^69,99^. Therefore, we hypothesized that EV formation, cargo selection, and release during necroptosis are highly regulated and selective, and exclusively characterize the necroptotic EVs compared with the control EVs.

To test this hypothesis and further profile the extracted EVs content, we performed principal component analysis between biological replicates. This analysis revealed that samples separate based on treatment (None vs. TBQ), supporting the biological interest in characterizing the necroptotic EV-specific cargo (Fig. 2A). Furthermore, unsupervised hierarchical clustering of 2,984 identified proteins separated samples into two distinct groups – the control and the necroptotic EVs (Fig. 2B). This indicates that the necroptotic EVs are characterized by a distinct proteome signature. Statistical analysis using a one-side Student’s t-test for paired samples yielded 352 proteins significantly upregulated in necroptotic vs. control EVs, with an FDR cutoff of 0.1 and S0 cutoff of 0.1, from hereon termed “TBQ-upregulated proteins” (Fig. 2C and online supplementary file “TBQ-upregulated proteins”). TBQ-upregulated proteins share 261 and 107 proteins with the exosome proteome data bases, Exocarta and Vesiclepedia, respectively, and contain 84 unique proteins (supplementary Fig. S3). Among these unique proteins are MLKL, the terminal executor of necroptosis, as well as the initiator caspase-8 and the ESCRTIII member, CHMP4B (Fig. 2D).

**Figure 2.**
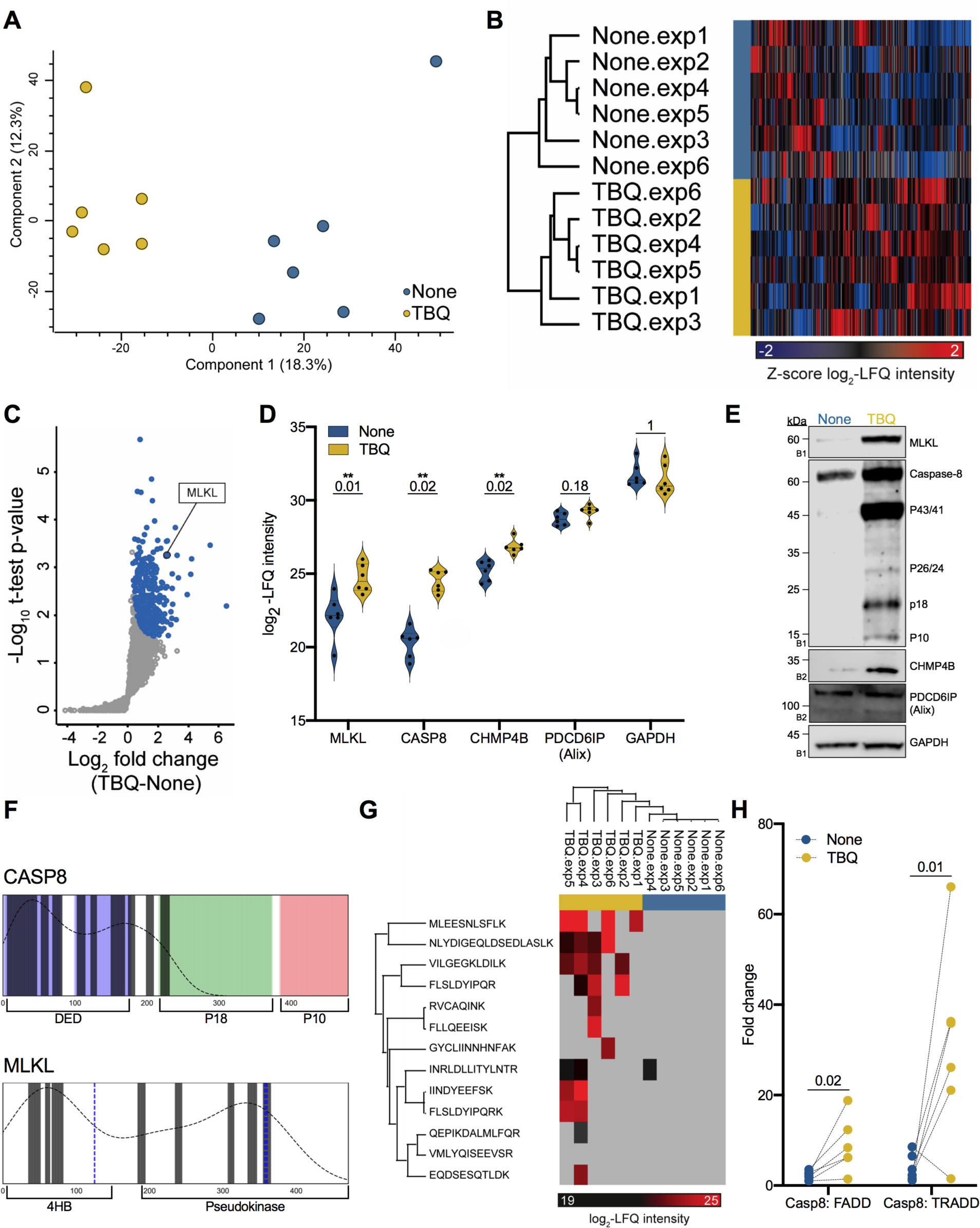
Proteome distinguishes between control and necroptotic EVs. **A**, Principal component analysis of biological replicates of extracted EVs showing that samples separate based on treatment (none vs. TBQ). **B**, Unsupervised hierarchical clustering analysis of 2,984 identified proteins between biological replicates, named by EV extraction. Color represents the Z-score of log_2_ -LFQ intensity of each protein. **C**, Volcano plot representing a one-side t-test analysis of necroptotic (TBQ) vs. control (none) EVs. 352 proteins were significantly upregulated in the necroptotic EVs and will be termed “TBQ-upregulated proteins” (blue, FDR cutoff=0.1, S0=0.1). MLKL is marked. **D**, Violin plots showing log_2_ -LFQ intensity of selected identified proteins. Q-value is mentioned individually above each plot, *Q < 0.1, **Q < 0.05. **E**, Validation of the selected EV proteins from D by immunoblot analysis of EVs from untreated and TBQ-treated cells using antibodies against MLKL, caspase-8, CHMP4B, PDCD6IP (Alix), and GAPDH. **F**, Alignment of the full-length caspase-8 and MLKL protein structures with peptides detected by MS analysis (marked as bars). Key structures of caspase-8 are colored and labeled: DED domains in blue, p18 subunit in green, and p10 subunit in red. Key structures of MLKL and labeled and phosphorylation sites are marked in blue. The dotted curve represents the density of the detected peptides along the full protein structure. **G**, Hierarchical clustering analysis of caspase-8 peptides detected by MS analysis. Color represents the log_2_ -LFQ intensity of each peptide, grey represents undetected levels. **H**, Dot plot showing the fold change in LFQ intensity between caspase-8 and FADD, or TRADD, paired by experiment. p-value for the multiple t-test is shown. **E**, EVs for validation were extracted independently of the six pairs of biological replicates that were analyzed by mass spectrometry. The number on the left of each blot indicates the blot number, each corresponding to an independent EV extraction. EVs, extracellular vesicles; TBQ, TNF-*α*, Birinapant and QVD-OPh; LFQ, label free quantification, CASP8, caspase-8; FDR, false discovery rate; 4HB, N-terminal four-helix bundle.

To validate these findings, we performed immunoblot analysis of these selected proteins from an independent EV extraction. We found that MLKL, caspase-8, and CHMP4B are indeed detected at higher levels in necroptotic EVs compared with control EVs, while the levels of the exosome marker, PDCD6IP (Alix), and GAPDH were unchanged, as expected given the proteomics data showed a non-significant fold change for both (Fig. 2E).

We further examined the specific caspase-8 and MLKL peptides that were detected in the extracted EVs by our MS analysis. While the detected MLKL peptides are distributed randomly along the protein, the capsase-8 peptides are almost entirely aligned with the death effector domain (DED)_1_-DED_2_ region, including the key residues involved in the DED-mediated filament assembly between death-inducing signaling complex (DISC) components^100^ (Fig. 2F). Quantitative analysis of the detected caspase-8 peptides revealed that the few peptides that are not aligned with the DED domains, *i.e*., EQDSESQTLDK and GYCLIINNHNFAK, were detected only in a single sample (Fig. 2G). This corresponds with our immunoblotting validation, showing high levels of the DED-containing caspase-8 forms, the full-length and p43/41, but low levels of the other cleavage products. We further assessed the presence of other DISC or TNF signaling complex components within the extracted EVs. While FADD and TRADD were detected, FLICE-like inhibitory protein (FLIP), RIPK1, and RIPK3 were not (supplementary table S2). The adaptors FADD and TRADD, both containing DED domains and capable of binding caspase-8 within the DISC, were not significantly upregulated in the necroptotic EVs. By comparing caspase-8 to the FADD and TRADD LFQ intensity in each sample, we emphasize that the ratio between these three proteins is not maintained among the control and the necroptotic EVs (Fig. 2H). This fits into the known model of TNF-*α* signaling.

In summary, necroptotic EVs are characterized by a unique proteome signature with 352 significantly upregulated proteins, as revealed by MS and supported by validation with immunoblotting.

### Components of ESCRTIII machinery and inflammatory signaling are enriched in the necroptotic EVs

To explore the distinctive signature of TBQ-upregulated proteins we performed gene ontology (GO) and Kyoto encyclopedia of genes and genomes (KEGG) pathways enrichment analysis (Fig. 3A, supplementary table S3 and online supplementary file “enrichment analysis”). We used the list of EV proteins from the Vescilepedia database^85,86^ as the background against which the TBQ-upregulated proteins were tested (Fisher’s exact test, Benjamini-Hochberg FDR < 0.02). Thus, the enriched processes represent the distinct signature of the necroptotic EVs in comparison to other EV systems, not to the general human proteome. As predicted, the “necroptosis” process was significantly enriched, supporting the specificity of the experimental system. Necroptosis here is defined as an inflammatory cell death pathway. As opposed to apoptotic cells, whose content is well-contained and rapidly cleared by phagocytosis, necroptotic cells profligately release cellular content as DAMPs following membrane permeabilization, triggering inflammation^2,5,32,33^. We surmised that the inflammatory potential of necroptosis may, in part, be due to the contribution of necroptotic EVs. In support of this, we found additional GO biological processes that are significantly enriched in necroptotic EVs include inflammatory signaling pathways, such as the “Toll-like receptor signaling pathway” and the “regulation of type I interferon production”. The ESCRT family is a group of proteins that facilitates protein transport via endosomes, multivesicular bodies, and vesicle budding^67^. The ESCRTIII members, CHMP2A and CHMP4B, were found to colocalize with pMLKL near the plasma membrane during necroptosis, thus suggesting a function in the release of necroptotic EVs^65,66,68,93^. In agreement, the cellular compartment annotated as “ESCRTIII” was enriched in necroptotic EVs. In addition, a group of proteins annotated as “phospholipid binding” was also enriched in necroptotic EVs. Among these are many members of the annexin family, *i.e*., annexin A1, A4, A6, A7, and A11, suggesting their involvement in the release of PS-exposing necroptotic EVs, and a potential mechanism for the recognition of necroptotic EVs by recipient cells.

**Figure 3.**
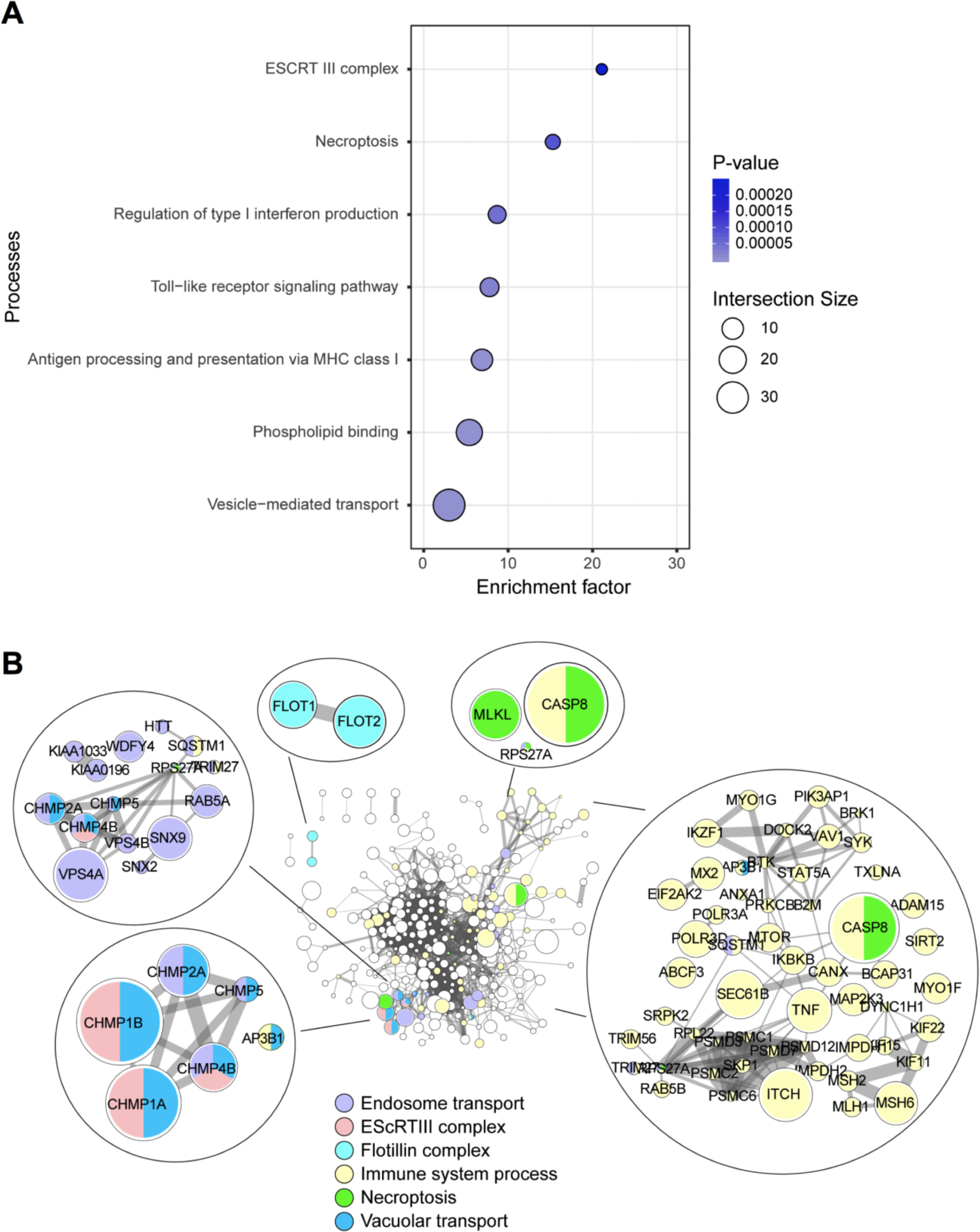
Enrichment and protein-protein interaction analyses reveal vesicle and inflammation pathways enriched in the necroptotic EVs. **A**, Processes enriched in the TBQ-upregulated proteins (Fisher exact test, FDR= 0.02, compared to the Vesiclepedia human proteome database). Color bars represent fisher-test p-values, symbol size represents intersection size between our data set and the GO category. **B**, Protein-protein interaction network of the TBQ-upregulated proteins (based on the String database). Node size represents fold change of TBQ vs. control. Edge width represents protein-protein co-expression level. Marked in color are proteins belonging to selected enriched biological processes. EVs, extracellular vesicles; TBQ, TNF-*α*, Birinapant and QVD-OPh; GO, gene ontology; FDR, false discovery rate.

Next, to get further insight into the biological events governing necroptotic EVs and their connectivity, we generated a protein-protein interaction network of the TBQ-upregulated proteins based on the String protein-protein interaction database. We found well-documented connections between proteins that function in vesicle formation, transport, and release, as well as in inflammation (Fig. 3B). As these connections are established in current literature, this suggests that EV formation and release during necroptosis is also a regulated and selective process.

Overall, this global profiling of the protein cargo of necroptotic EVs indicates possible mechanisms for their upstream formation and release, and their downstream binding, uptake, and inflammatory effects on recipient cells. Hence, this data analysis is the first step towards uncovering new players in necroptotic signaling, EV biogenesis, and their biological effects.

### Necroptotic EVs contain new unstudied components of vesicle formation and transport, and necroptosis signaling pathways

As mentioned above, we^64^ and others (Yoon *et al*.^66^ and Gong *et al*.^65^), have previously demonstrated PS exposure during necroptosis. We reported that PS-exposing necroptotic U937 cells release EVs early during necroptosis, which contain pMLKL^64^. Yoon *et al*. have shown that MLKL controls the formation of intraluminal and EVs, and that its activation during necroptosis enhances the release of EVs^66^. In addition, they have studied the proteins that are specifically associated with MLKL within EVs by immunoprecipitating MLKL from lysed EVs extracted from TNF-*α*, Birinapant and z-VAD-fmk (TBZ)-treated HT-29 cells. In light of these results, we compared our TBQ-upregulated proteins with the major proteins reported to precipitate with MLKL by Yoon *et al*. (Fig. 4A). We found eight proteins in common, including the ESCRTIII members, CHMP1A, CHMP1B, CHMP2A, CHMP5, and IST-1. The lipid raft-associated proteins, flotillin-1 and flotillin-2^101–103^, were also found by both analyses (Fig. 4B). This suggests that lipid rafts, which are microdomains of the plasma membrane abundant with cholesterol and glycosphingolipids^104,105^, are involved in the release of necroptotic EVs. Interestingly, it has recently been shown that flotillin-1 and flotillin-2 delay necroptosis by removing pMLKL from membranes, via flotillin-mediated endocytosis followed by localization to lysosomes and degradation^106^. Their presence within the necroptotic EVs, confirmed by both our and Yoon *et al*.’s systems, suggests that this mechanism might not solely culminate in MLKL degradation but also in its exocytosis.

**Figure 4.**
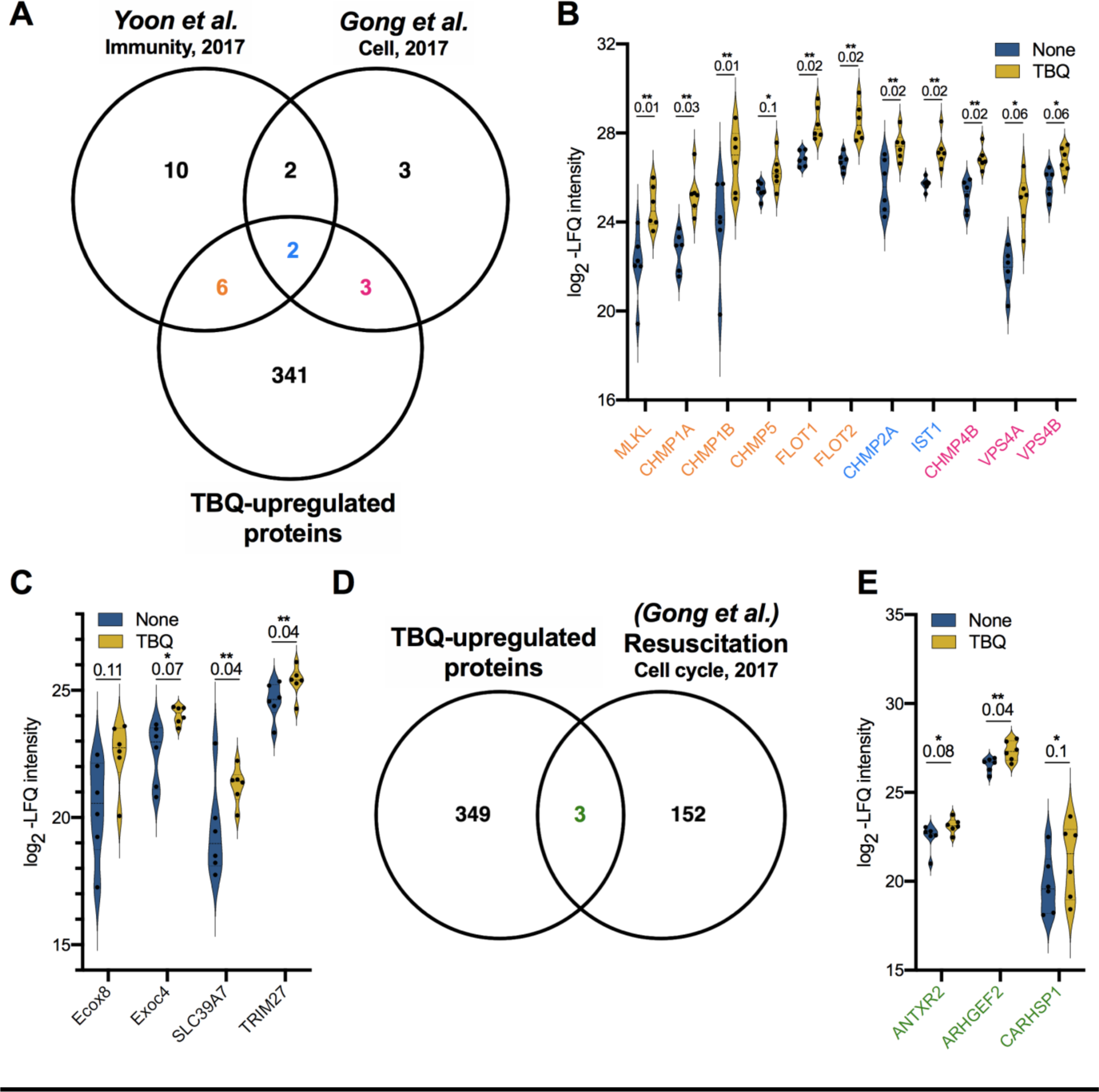
Necroptotic EVs are abundant with reported and unreported components of vesicle formation and transport and necroptosis signaling pathways. **A**, Venn diagram of the TBQ-upregulated proteins compared with proteins that were found using mass spectrometry to associate with MLKL within EVs by *Yoon et al*., Immunity, 2017^66^ and with proteins that were shown to regulate EV formation and sensitivity to necroptosis by *Gong et al*., Cell, 2017^65^. **B**, Violin plots showing log_2_ -LFQ intensity of overlapping proteins. Protein names are colored according to their overlapping groups in A. Q-value is mentioned individually above each plot, *Q < 0.1, **Q < 0.05. **C**, Violin plots showing log_2_ -LFQ intensity of selected proteins that were upregulated in necroptotic (TBQ) vs. control (none) EVs. Q-value is mentioned individually above each plot, *Q < 0.1, **Q < 0.05. **D**, Venn diagram of the TBQ-upregulated proteins compared with genes that were found by RNA-Seq to be upregulated during resuscitation from necroptosis by *Gong et al*., Cell Cycle, 2017^68^. **E**, Violin plots showing log_2_ -LFQ intensity of overlapping proteins from D (green). Q-value is mentioned individually above each plot, *Q < 0.1, **Q < 0.05. EVs, extracellular vesicles; TBQ, TNF-*α*, Birinapant and QVD-OPh; FDR, false discovery rate.

Gong *et al*. revealed that the ESCRTIII complex acts downstream of pMLKL to induce the shedding of MLKL-damaged membrane, consequently delaying cell death execution^65^. By silencing many ESCRT family members and related accessory proteins in necroptotic cells, they found 10 members of the family for which silencing resulted in reduced shedding of membrane bubbles, increased sensitivity to necroptosis, or a reduced ability to recover from early necroptosis upon stimulant removal (termed “resuscitation”). Five of these proteins were also significantly upregulated in our necroptotic EVs, and two of them were also found in the analysis by Yoon *et al*. (Fig. 4A and Fig. 4B).

Beyond the overlap with established necroptosis systems, we looked for other proteins yet unstudied in this context. Using a less stringent cutoff, we extended our analysis and searched for additional biologically-relevant proteins that were upregulated in necroptotic EVs. This revealed proteins such as exocyst complex component-8 (Exoc8, also known as Exo84), Exoc4 (also known as Sec8), solute carrier family 39 member 7 (SLC39A7), and tripartite motif containing 27 (TRIM27) (Fig. 4C). Exoc8 and Exoc4 are components of the exocyst complex that tethers vesicles to the plasma membrane, resulting in their soluble M-ethylmaleimide-sensitive factor attachment protein receptors (SNAREs)-mediated fusion and secretion^107,108^. Of note, some SNARE proteins were identified in the necroptotic EVs (*i.e*., were detectable in at least three of the six TBQ samples), although not significantly upregulated when compared with control EVs (supplementary table S4). Moreover, the exocyst complex binds Rab proteins on vesicles and is considered a Rab effector^109–111^. Rabs are small GTPase RAS-related proteins, and in their GTP-bound form they facilitate the recruitment of tethers to specific cellular locations^108,112^. Several Rab proteins were identified in the necroptotic EVs (supplementary table S5), and among those significantly upregulated were RAB5A, RAB5B, RAB5C, and RABGAP1. In support, Yoon *et al*. have shown that RNA silencing of the Rab proteins, RAB27A and RAB27B, in HT-29 necroptotic cells and LPS-treated caspase-8-deficient bone-marrow-derived dendritic cells (BMDCs) results in a reduced number of EVs and increased cell death^66^. Thus, these mechanisms may function in necroptotic EV biogenesis.

The zinc transporter, SLC39A7, which was significantly upregulated in the necroptotic EVs, was found to be needed for TNF-*α*-induced necroptosis in KBM7 cells by modulating TNFR1 trafficking from the endoplasmic reticulum (ER) to the cell surface^113^. *SLC39A7*^-/-^ cells demonstrated impaired ER homeostasis. TRIM27, which we also found to be significantly upregulated in the necroptotic EVs, positively regulates TNF-*α*-induced apoptosis^114^. TRIM27 induces ubiquitin-specific-processing protease 7 (USP7)-mediated RIPK1 de-ubiquitination, resulting in apoptosis upon stimulation. Thus, TRIM27 might be similarly required for necroptosis when capsase-8 is inhibited. These findings suggest that the association between necroptotic EV release and the delayed execution of cell death might be also attributed to new unstudied proteins, rather than the shedding of membrane-damaging MLKL alone.

We, and others, have previously shown that PS-exposing necroptotic cells are not committed to die^64,65,68^. By the addition of the pMLKL-translocation inhibitor, necrosulfonamide (NSA), we were able to rescue sorted PS-positive U937 necroptotic cells. Gong *et al*. suggested that necroptotic cells are rescued by ESCRTIII-mediated shedding of pMLKL-incorporated damaged membranes. By utilizing their dimerizable RIPK3 or MLKL system, they analyzed gene expression in the resuscitated necroptotic cells compared with untreated controls, using RNA sequencing. To test the hypothesis that necroptotic EV release delays cell death, we compared the TBQ-upregulated proteins with the genes that were found to be upregulated during necroptosis resuscitation (Fig. 4D and Fig. 4E). Interestingly, we found that mRNA transcripts from only three proteins were upregulated in the resuscitated necroptotic cells. One of the three proteins is calcium-regulated heat-stable protein 1 (CARHSP1), a cytoplasmic protein that localizes to exosomes to increase TNF-*α* production by enhancing TNF-*α* mRNA stability^112^. This result suggests that proteins with a key role in necroptosis are secreted in EVs, while resuscitation-supporting proteins are not. However, this finding might also be influenced by many other differences between the two systems.

Altogether, our results show that necroptotic EVs carry cargo involved in many relevant pathways, but also contain numerous proteins that have not yet been studied in this context. Further investigation of these proteins could lead to the discovery of new genetic and pharmacological approaches to regulate necroptosis, including necroptotic EV formation, release, uptake, and their impact on the microenvironment.

### Necroptotic EVs are phagocytosed to promote inflammation in recipient cells

We previously reported that PS-exposing necroptotic cells are phagocytosed and trigger the secretion of TNF-*α* and the potent monocyte-attracting chemokine, CCL2 (MCP-1), at higher levels than following the uptake of early apoptotic cells^64^. To assess whether necroptotic EVs share this potential, we utilized our established system to examine uptake of necroptotic EVs by thioglycolate-derived peritoneal macrophages (TG-macs) and its effect (Fig. 5A). While EVs were phagocytosed by TG-macs as efficiently as control and apoptotic EVs (Fig. 5B), they induced an increased secretion of the pro-inflammatory cytokines IL-6 and TNF-*α*, as well as the chemokine CCL2^115–117^ (Fig. 5C). These results suggest that necroptotic EVs can be taken up by recipient cells, promoting inflammatory signaling.

**Figure 5.**
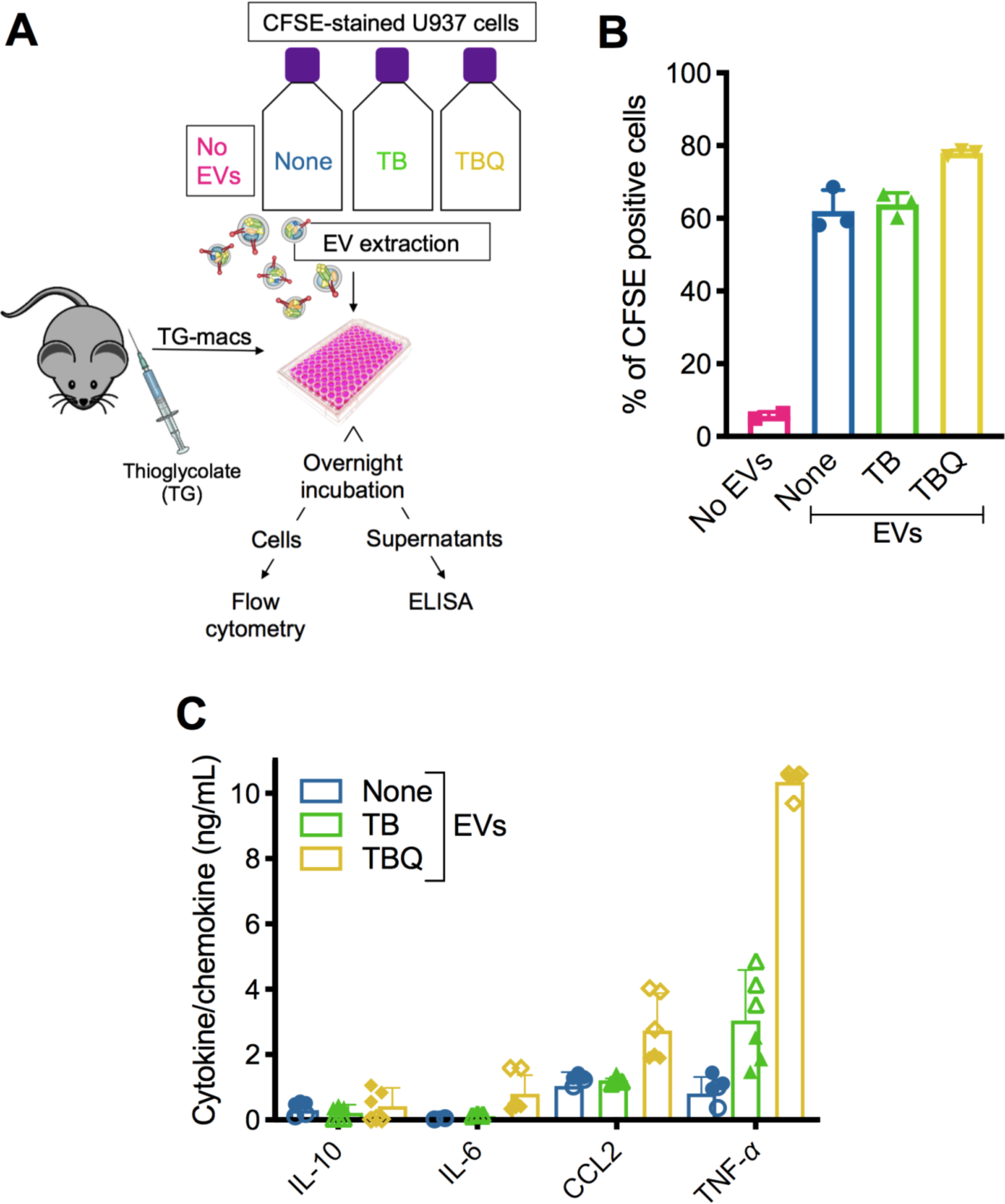
Necroptotic EVs modulate cytokine and chemokine secretion. **A**, Schematic overview of EV phagocytosis assay. EVs from untreated (none), apoptotic (TB), or necroptotic (TBQ) CFSE-stained U937 cells were added to thioglycolate-derived peritoneal macrophages (TG-macs) overnight. **B**, TG-macs were then harvested and analyzed by flow cytometry for their CFSE-positivity, demonstrating uptake of the EVs. **C**, Supernatants were collected and secreted cytokine and chemokine levels were analyzed by ELISA. **B**, Data are presented as the mean of triplicate samples ± SD. **C**, Empty symbols are triplicate samples of mouse#1, colored symbols are triplicate samples of mouse#2. Data are presented as the mean of duplicate mice ± SD and are representative of two independent experiment, n = 5. TG-macs, thioglycolate-derived peritoneal macrophages; EVs, extracellular vesicles; T, TNF-*α*; TB, TNF-*α* and Birinapant; TBQ, TNF-*α*, Birinapant and QVD-OPh; PI, propidium iodide; h, hours; SD, standard deviation.

### Necroptotic EVs carry cancer neoantigens

Recently, necroptosis has been suggested as a potential therapeutic mechanism to overcome the ability of tumor cells to escape apoptosis^118,119^. Thus, instead of using anti-cancer drugs (chemotherapeutics) that trigger apoptosis, drugs that induce necroptosis might be better suited to specifically kill tumor cells. In parallel, anti-cancer immunotherapy has emerged as an effective therapeutic approach in a number of cancers, educating the immune system to recognize and eliminate tumor cells. The inflammatory nature of necroptosis has been utilized to this end by few groups, who have reported that administration of necroptotic cancer cells drives anti-tumorigenic adaptive immune responses^54–58^. The exact mechanism of this effect is not yet fully understood. In this regard, proteins delivered by exosomes secreted from intestinal epithelial cells or dendritic cells are internalized into recipient dendritic cells, processed, and presented as antigens, potentially triggering an immune response^120,121^. Thus, we hypothesized that necroptosis has the potential not only to kill tumor cells, but also to deliver tumor neoantigens via necroptotic EVs to activate an anti-tumorigenic adaptive immune response. To address this hypothesis, we compared all the proteins identified in the necroptotic EVs with neoantigens in a dataset derived from B16F10, CT26, 4T1, or CT26 cell lines or in an *in silico* prediction model, which contains immunogenic antigens recognized by CD4^+^ T cells that elicit an anti-tumor response^122^. We found a total of 26 potential neoantigens in our necroptotic EVs, with six of them being significantly, or almost significantly, upregulated in comparison to control EVs (Fig. 6A).

**Figure 6.**
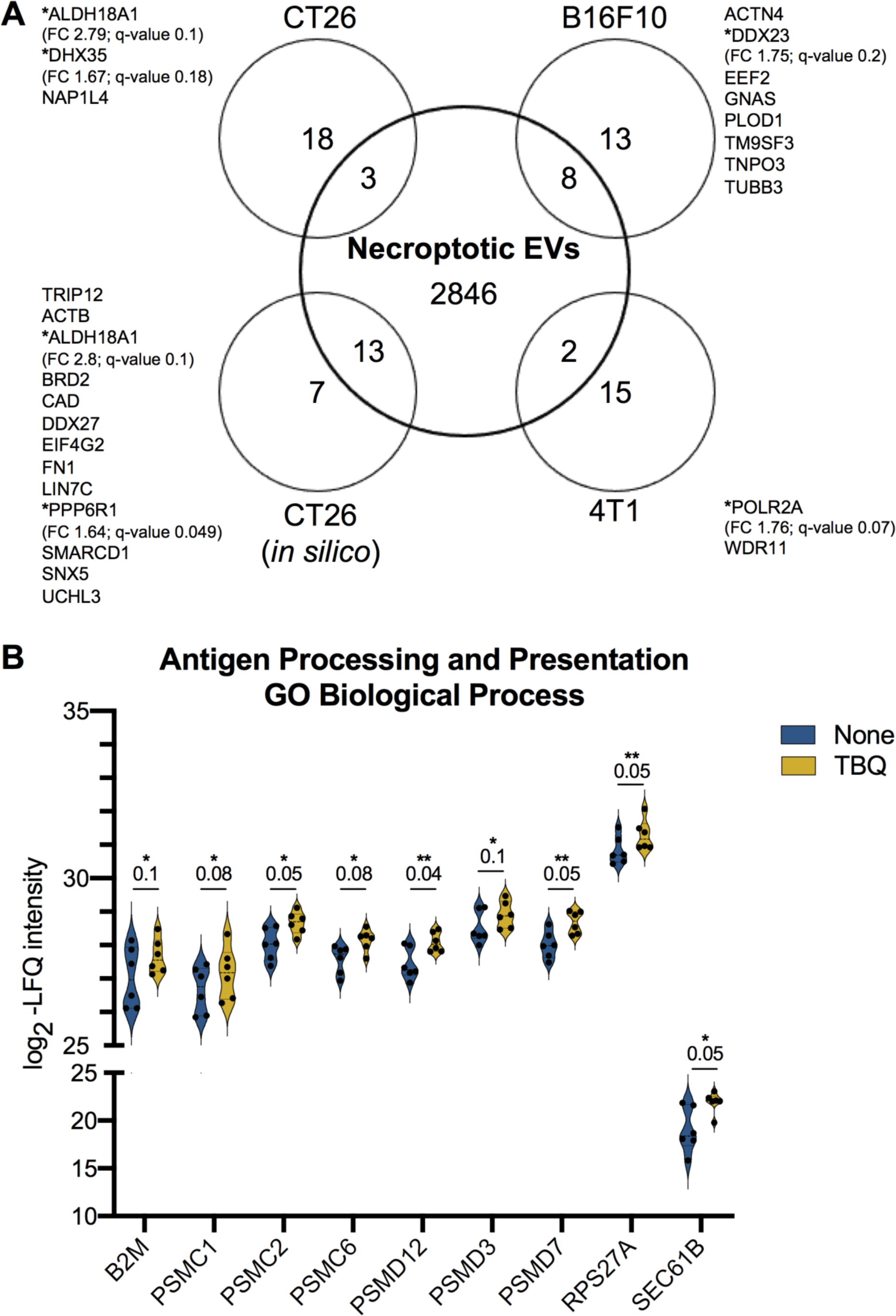
Necroptotic EVs contain tumor neoantigens. **A**, Venn diagram of total proteins identified in necroptotic EVs compared with the published tumor neoantigens from B16F10, CT26, 4T1 models or CT26 *in silico* prediction^122^. TBQ-upregulated proteins with FDR cutoff of 0.2 are marked with **\***; Fold change (FC) and q-value are mentioned for each. **B**, Violin plots showing log_2_ -LFQ intensity of the TBQ-upregulated proteins annotated as the enriched GO biological process “antigen processing and presentation via MHC class I”. Q-value is mentioned individually above each plot, *Q < 0.1, **Q < 0.05. EVs, extracellular vesicles; FDR, false discovery rate; FC, fold change; TBQ, TNF-*α*, Birinapant and QVD-OPh; GO, gene ontology; LFQ, label free quantification; MHC, major histocompatibility complex.

Mast cell- and macrophage-derived EVs contain signals, such as heat shock protein 60 (HSP60), heat shock protein family A member 8 (HSPA8), and pathogen-associated molecular patterns (PAMPs), that stimulate activation of recipient cells into immunogenic APCs^123–125^. In support, our analysis found that the GO biological process annotated “antigen processing and presentation of exogenous peptide antigen via major histocompatibility complex (MHC) class I” was significantly enriched in the TBQ-upregulated proteins (Fig. 3A), and included proteins, such as *β*2-microglobulin (B2M), proteasome subunits, and SEC61B (Fig. 6B), suggesting that necroptotic EVs can facilitate activation and antigen presentation by APCs upon uptake.

In summary, these results suggest that necroptotic EVs could serve as a potential delivery system for tumor neoantigens, promoting their effective presentation by APCs.

## Discussion

Although necroptosis has been known for two decades as a regulated and inflammatory form of cell death^2,126^, neither its regulation nor inflammation-inducing consequences have been fully elucidated. We, and others, have recently reported that PS exposure on the outer cell membrane is not exclusive to apoptosis and its immunologically silent nature^92^, but also occurs during early necroptosis, prior to plasma membrane permeabilization^64–66,93,94^. We have also shown that necroptotic cells release PS-exposing EVs that are abundant in proteins, but have a low-nucleic acid content in comparison to apoptotic bodies.

Here, we uncover the proteome of necroptotic EVs using mass spectrometry-based proteomics of ultracentrifugation-extracted EVs from necroptotic and untreated U937 cells. In this study we show that necroptotic EVs have a unique proteome signature with significant upregulation of many proteins, *e.g*., MLKL and caspase-8, which were also validated by immunoblotting analysis.

Intrigued by the presence of caspase-8 in the necroptotic EVs, we examined the specific peptides that were detected. Caspase-8 is an initiator caspase that is recruited to intracellular multi-protein complexes, such as the DISC or TNF signaling Complex I and II, following death receptor ligation^22,100,127^. This assembly is mediated by binding of DED domains shared by caspase-8, FADD, and TRADD, resulting in homo-oligomerization, cleavage, and activation of caspase-8 and subsequent apoptosis, or necroptosis when capsase-8 activity is inhibited^8,10,11,128–130^. Kinetics studies of procaspase-8 cleavage at the DISC suggest that the first cleavage event is at position D384, resulting in the formation of the p10 and p43/41 fragments, the latter of which is subsequently cleaved at position S216 to generate the p18 and p26/24 subunits^131^. Treatment with QVD-OPh was shown to only partially block the first cleavage event, but to fully prevent the second cleavage event^132^. In agreement, we detected mainly full-length and p43/41 caspase-8 products by immunoblot and peptides that are almost exclusively aligned with the DED domain by MS. This suggests that the p10 product remains within the necroptotic cells, while the full-length and p43/41 products are selectively secreted in necroptotic EVs. Of note, it is also possible that the second cleavage event does occur to some extent (as we detect low levels of p18 by immunoblotting), but that the remaining p26/24 is not detectable due to the specificity of the caspase-8 antibody and the detection limit of MS analysis. Following DISC formation, both the p43/41 and p26/24 subunits, which contain the DED domains, remain bound to, and contained within, the DISC^133,134^. This corresponds with the fact that the DED-containing adaptors, FADD and TRADD, were also found in the necroptotic EVs. The fact that caspase-8 was significantly upregulated in the necroptotic EVs, while FADD and TRADD were not, matches the known stoichiometric ratios between caspase-8 and these adaptors within the DISC, as one adaptor molecule recruits many caspase-8 molecules to homo-oligomerize^131,135^. In addition, the presence of the lipid raft-associated proteins, flotillin-1 and flotillin-2, supports the release of these membrane-bound DISC components. Notably, FLIP, RIPK1, and RIPK3 were not detected, suggesting that TNF signaling Complex II is absent from the necroptotic EVs, but this result might also stem from MS detection limits. In addition, the absence of the death receptors might be due to the ectodomain shedding reported to be involved in necroptosis^136,137^.

We found that components of ESCRTIII machinery and inflammatory signaling are enriched in necroptotic EVs. Gong *et al*. first reported that PS exposure during necroptosis is mediated by ESCRTIII machinery, which also induces the budding of pMLKL-containing damaged membranes to delay the execution of cell death^65^. We have previously speculated whether the same mechanism is involved in the formation of necroptotic EVs^64^. Interestingly, we found that many ESCRTIII members are upregulated in necroptotic EVs, some of which were shown by Gong *et al*. to play a role in decreasing sensitivity to necroptosis and enhancing recovery from early necroptosis. This is a key finding, bridging the system of Gong *et al*. and our previous report of EV release during necroptosis.

Among the enriched processes in necroptotic EVs we also found the molecular function group “phospholipid binding” that includes many annexin family members. Annexins bind phospholipids in a calcium-dependent manner. Due to this ability, they are localized to membrane domains where they can pull and fuse two membranes together, thus functioning in exocytosis and vesicle secretion in response to calcium sensing^138^. Thus, it is not surprising that annexins are among the cytosolic proteins that should be analyzed to demonstrate the nature and purity of EVs^139^. Annexins are suggested to have a role in vesicle docking to recipient cells, by binding an unknown tether on the recipient plasma membrane next to an adjacent binding of PS to its receptor^69,140^. Therefore, our finding proposes the involvement of annexins in both the release of PS-exposing necroptotic EVs and their recognition by recipient cells.

Moreover, by generating a protein-protein interaction network for the cargo of necroptotic EVs, we found established connections between proteins that function in vesicle formation, transport, release, and in inflammatory signaling. This identifies necroptotic EV formation and cargo assortment as regulated and selective processes.

Our results strongly imply there are still unknown connections to be established as necroptotic EV research develops. We found that flotillin-1 and flotillin-2, which are lipid raft-associated proteins, are significantly upregulated in necroptotic EVs, suggesting that lipid rafts play a role in the release of necroptotic EVs. In agreement, lipid-rafts are involved in PS-exposing EV release from activated platelets, monocytes, endothelial cells, and erythrocytes^141–144^. This involvement is thought to be calcium-dependent in many cases, but this is mostly unclear.

Transmembrane protein 16F (TMEM16F) is a calcium-dependent phospholipid scramblase^93,145,146^. Following pore formation by bacterial toxins and other agents, TMEM16F was recently shown to be essential for PS exposure and plasma membrane repair, via the release of EVs^147^, supporting calcium efflux as the bridge between PS exposure and EV release during necroptosis. In support, our results point to the involvement of several calcium-dependent vesicle-transport machineries, evidenced by the upregulated presence of annexins, Rabs, SNAREs, exocyst complex proteins, ESCRTIII members, and lipid rafts within necroptotic EVs. While TMEM16F (ANO6) was detected in both necroptotic and control EVs, it was not significantly upregulated within necroptotic EVs.

The presence of exocyst complex members, SNAREs, and Rab proteins in necroptotic EVs was surprising. Inward budding of early endosomal membrane forms ILVs that incorporate to form MVEs. Following MVE formation and transport, the final step of exosome secretion is MVE fusion to the plasma membrane, resulting in ILV release as exosomes^69^. This step is mediated in some cells by Rabs and their effectors from the exocyst complex, tethering vesicles to plasma membrane and inducing SNARE-mediated secretion^107–112,148–150^. So far, these mechanisms have not been implicated in necroptosis but should now be studied in this context in light of the upregulation of exocyst components in necroptotic EVs. We found many other, as yet, unreported proteins with the potential to be involved in necroptotic EV biology and necroptotic signaling, such as SLC39A and TRIM27.

Although necroptosis is considered to be a highly inflammatory form of cell death^2,151^, it has previously been shown to attenuate TNF-*α*- or lipopolysaccharide (LPS)-induced inflammation, by terminating the production of multiple cytokines and chemokines^152^. In support, necroptotic cells demonstrate reduced release of conventionally-secreted cytokines, such as CCL2 and IL-8, by MS and ELISA in comparison to apoptotic cells^136^. However, these reports do not take into account the inflammatory impact necroptotic cells have on recipient cells in the microenvironment. We have previously shown that PS-exposing necroptotic cells are phagocytosed and trigger the secretion of TNF-*α* and CCL2 at higher levels compared to the uptake of early apoptotic cells^64^. Here we report, for the first time to our knowledge, that necroptotic EVs can similarly be phagocytosed by macrophages, triggering an enhanced secretion of cytokines and chemokines in comparison to apoptotic EVs. Hence, this discrepancy can be also settled by the influence of necroptotic EVs on their environment. In addition, necroptotic EVs appear to mediate a delay in cell death execution, which is thought to be ESCRTIII-dependent, as well as sustain cytokine production and secretion from necroptotic cells and support necroptosis-induced inflammation^93^. Finally, we have discovered that necroptotic EVs contain tumor neoantigens that are recognized by T cells, and may promote antigen presentation to support an anti-tumor adaptive immune response. As vaccination with necroptotic cancer cells was shown to facilitate anti-tumor immunogenicity in mice^54^, necroptotic EVs might contribute, in part, to this phenomenon.

Of note, the use of a caspase-inhibitor to induce necroptosis might interfere with meaningful proteolytic events during this process. In addition, post-translational modifications (PTM), such as phosphorylation, might be missed by the bottom-up MS sample processing approach^97^. Thus, future research should aim to use a necroptotic stimulus that does not include pharmacological caspase inhibition, and also to apply alternative proteomic approaches, *e.g*., PTM proteomics or semi-tryptic digest of proteins. In addition, future studies should investigate EV cargo other than proteins, such as mRNA and miRNAs.

To conclude, we show that necroptotic cells release EVs via specific pathways that are yet to be fully characterized, and that these EVs can affect the microenvironment (Fig. 7). Our study establishes necroptotic EV release as a highly-regulated, selective, and effective phenomenon, with the potential to shed light on cell-fate-determining players and inflammation-promoting necroptotic mechanisms. Finally, we suggest that necroptotic EVs may ultimately be harnessed for cancer vaccination.

**Figure 7.**
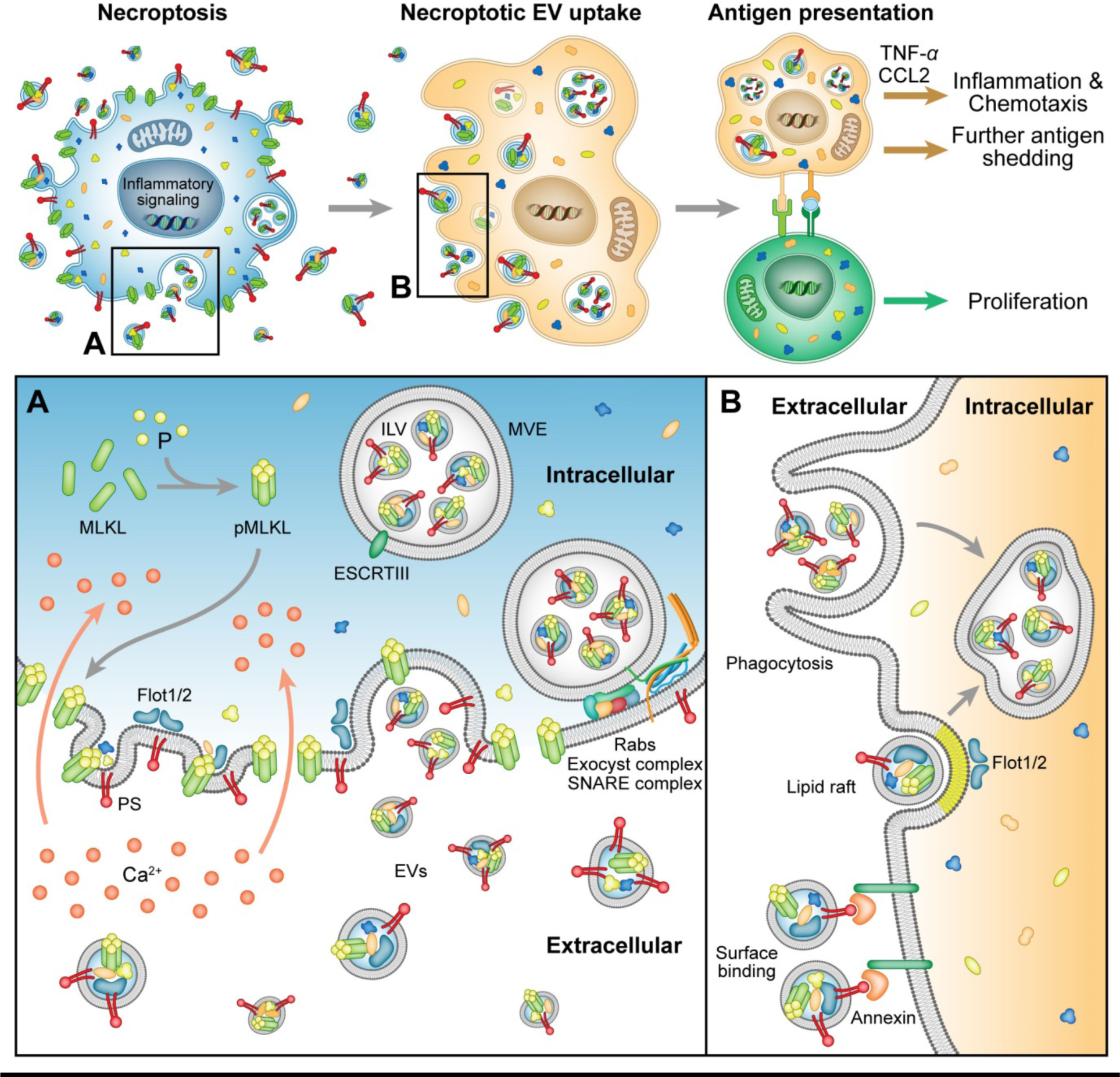
Suggested model of necroptotic EV release, uptake, and effects. Necroptotic cells expose PS on the outer plasma membrane and release EVs via specific pathways, involving pMLKL-mediated Ca^2+^ influx, ESCRTIII machinery, Rabs, SNAREs, exocyst complex, and lipid rafts. EVs are then taken up by recipient cells via phagocytosis, membrane fusion, or binding of PS and annexin, resulting in inflammation, chemotaxis, and antigen presentation to T cells. EVs, extracellular vesicles; PS, phosphatidylserine; pMLKL, phosphorylated MLKL; ILV, intraluminal vesicle; MVE, multivesicular endosome.

## Acknowledgement

This work was performed in partial fulfillment of the requirements for a PhD degree (IS, GYA, ZE, SZ and HC), Sackler Faculty of Medicine, Tel Aviv University, Israel. We thank The Smoler Proteomics Center at the Technion for performing the mass spectrometry. We would also like to thank Dr. Mary Speir for editing this manuscript.

## Funding Details

This work was supported by the Recanati Foundation (TAU), received by MG; Israel Science Foundation (ISF) under Grants #1416/15 and 818/18, received by MG; Alpha-1 foundation under Grant #615533, received by MG; U.S. – Israel Binational Science Foundation (BSF) under Grant #2017176, received by MG; Individual research grant from Varda and Boaz Dotal Research Center in Hemato-Oncology, received by MG; Israel Science Foundation (ISF) under Grant #2235/16, received by NRR and Enid Barden and Aaron J. Jade President’s Development Chair for New Scientists in Memory of Cantor John Y. Jade, received by NRR. The funders had no role in study design, data collection and analysis, decision to publish, or preparation of manuscript.

## Declaration of Interest Statement

The authors declare no conflict of interest.

## Authors’ contribution

Conceptualization: IS, GYA, MG; Funding acquisition: MG, NRR; Investigation: IS, GYA, ZE, LEB, SZ, HC, YOB, NRR, MG; Methodology: IS, GYA, ZE, LEB, SZ, HC, MG; Supervision: LEB, MG; Writing - original draft: IS; Writing – review & editing: IS, GYA, ZE, LEB, HC, YOB, NRR, MG.

## Abbreviations

4HB: N-terminal four-helix bundle
A5: AnnexinV
APCs: antigen-presenting cells
B2M: *β*2-microglobulin
BMDC: bone-marrow-derived dendritic cells
CARHSP1: calcium-regulated heat-stable protein 1
CASP8: caspase-8
CFSE: carboxyfluorescein succinimidyl ester
CHMP2A: charged multivesicular body protein 2A
cIAPs: cellular inhibitor of apoptosis proteins
CLTC: Clathrin heavy chain 1
CMV: cytomegalovirus
DAMPs: danger associated molecular patterns
DED: death effector domain
DISC: Death-inducing signaling complex
EBV: Epstein-Barr virus
EDTA: ethylenediamine tetra-acetic acid
ER: endoplasmic reticulum
ESCRT: endosomal sorting complexes required for transport
EVs: extracellular vesicles
Exoc: exocyst complex component
FADD: FAS-associated death domain protein
FBS: fetal bovine serum
FC: fold change
FDR: false discovery rate
FLIP: FLICE-like inhibitory protein
FLOT: flotillin
GAPDH: glyceraldehyde 3-phosphate dehydrogenase
GO: gene ontology
h: hours
HCD: high energy collision dissociation
HMGB1: high mobility group box 1
HRP: horseradish peroxidase
HSP60: heat shock protein 60
HSPA8: heat shock protein family A member 8
ILVs: intraluminal vesicles
KEGG: Kyoto encyclopedia of genes and genomes
LC-MS/MS: liquid chromatography with tandem mass spectrometry
LFQ: label free quantification
LPS: lipopolysaccharide
MHC: major histocompatibility complex
MLKL: mixed lineage kinase domain-like
MVE: multivesicular endosome
NF-κB: nuclear factor kappa-light-chain enhancer of activated B cells
NSA: necrosulfonamide
NTA: nanoparticle tracking analysis
PAMPs: pathogen-associated molecular patterns
PCD6IP: programmed cell death 6-interacting protein
PI: propidium iodide
pMLKL: phosphorylated MLKL
PS: phosphatidylserine
PSM: peptide-spectrum match
PTM: post-translational modifications
RIPK1: receptor-interacting serine/threonine-protein kinase 1
RIPK3: receptor-interacting serine/threonine-protein kinase 3
SD: standard deviation
SDS-PAGE: sodium dodecyl sulphate-polyacrylamide gel electrophoresis
SEM: standard error of the mean; scanned electron microscopy
SLC39A7: solute carrier family 39 member 7
SMAC: second mitochondria-derived activator of caspases
SNAREs: soluble M-ethylmaleimide-sensitive factor attachment protein receptors
TB: TNF-*α* and birinapant
TBQ: TNF-*α*, Birinapant and QVD-OPh
TBZ: TNF-*α*, Birinapant and z-VAD-fmk
TG-macs: thioglycolate-derived peritoneal macrophages
TLRs: Toll-like receptors
TMEM16F: Transmembrane protein 16F
TNF-*α*: tumor necrosis factor-*α*
TNFR1: tumor necrosis factor receptor 1
TRADD: Tumor necrosis factor receptor type 1-associated death domain
TRAF2: TNF receptor associated factor 2
TRIM27: tripartite motif containing 27
TSG101: tumor susceptibility gene 101
TSZ: TNF-*α*, SMAC and z-VAD-fmk
USP7: ubiquitin-specific-processing protease 7

## Supplementary

**Figure S1.**
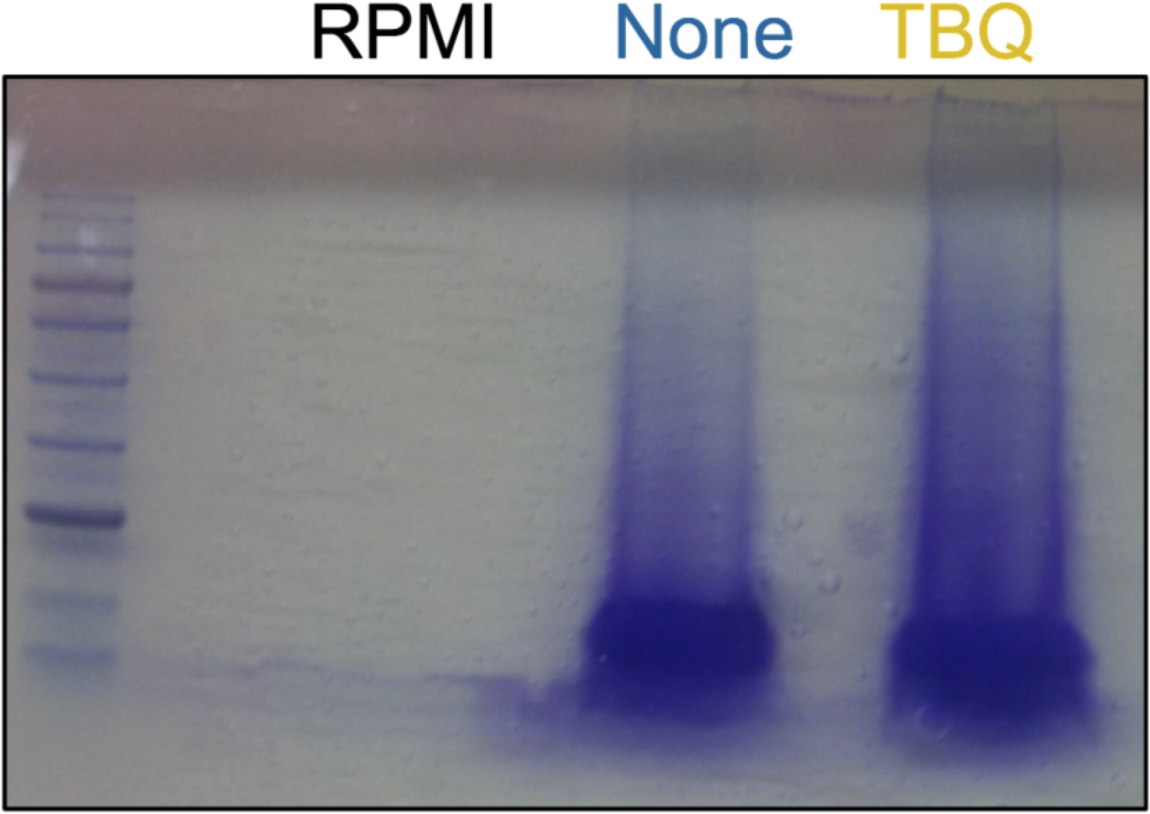
Coomassie blue staining of the control and the necroptotic EV.

**Figure S2.**
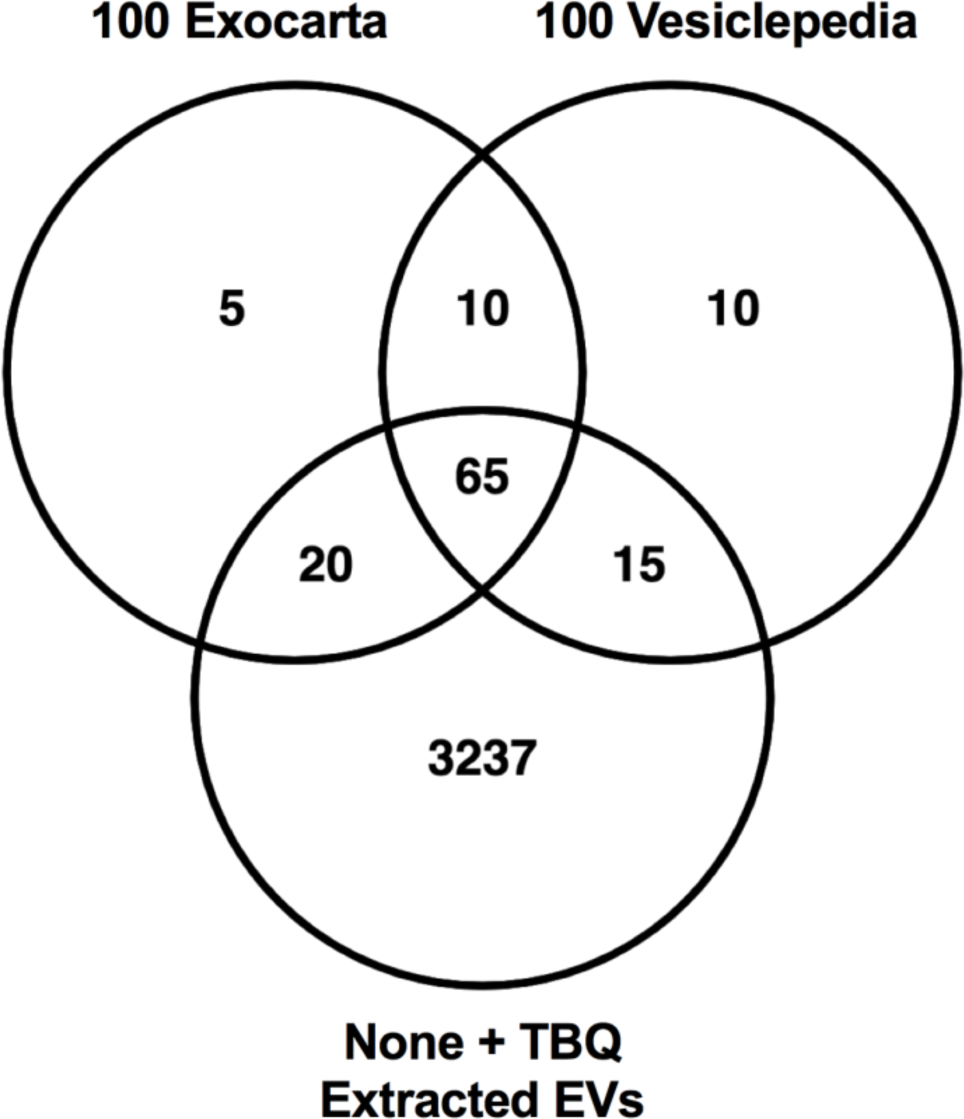
(Related to Fig. 1) Extracted EVs contain 65 of the 75 most frequently identified proteins in both Exocarta and Vesiclepedia. Venn diagram of total proteins identified in extracted EVs compared with the 100 most frequently identified proteins in Exocarta and and Vesiclepedia.

**Table S1.** (Related to Fig. 1) Common exosomal marker proteins identified in the extracted EVs.

| Gene Name | Protein Name | Number out of 100 most identified in Exocarta | Number out of 100 most identified in Vesiclepedia |
| --- | --- | --- | --- |
| PDCD6IP | Programmed cell death 6-interacting protein | 2 | 1 |
| HSPA8 | Heat shock cognate 71 kDa protein | 3 | 3 |
| GAPDH | Glyceraldehyde-3-phosphate dehydrogenase | 4 | 2 |
| ACTB | Actin, cytoplasmic 1 | 5 | 4 |
| CD63 | CD63 antigen | 7 | 12 |
| ENO1 | Alpha-enolase | 9 | 9 |
| HSP90AA1 | Heat shock protein HSP 90-alpha | 10 | 8 |
| TSG101 | Tumor susceptibility gene 101 protein | 11 | 39 |
| PKM | Pyruvate kinase | 12 | 7 |
| LDHA | L-lactate dehydrogenase A chain | 13 | 45 |
| YWHAZ | 14-3-3 protein zeta/delta | 15 | 13 |
| PGK1 | Phosphoglycerate kinase 1 | 16 | 16 |
| EEF2 | Elongation factor 2 | 17 | 21 |
| ALDOA | Fructose-bisphosphate aldolase A | 18 | 20 |
| HSP90AB1 | Heat shock protein HSP 90-beta | 19 | 11 |
| ANXA5 | Annexin A5;Annexin | 20 | 10 |
| FASN | Fatty acid synthase | 21 | 46 |
| YWHAE | 14-3-3 protein epsilon | 22 | 14 |
| CLTC | Clathrin heavy chain 1 | 23 | 17 |
| VCP | Transitional endoplasmic reticulum ATPase | 26 | 24 |
| TPI1 | Triosephosphate isomerase | 27 | 23 |
| PPIA | Peptidyl-prolyl cis-trans isomerase A | 28 | 18 |
| MSN | Moesin | 29 | 26 |
| CFL1 | Cofilin-1 | 30 | 25 |
| PRDX1 | Peroxiredoxin-1 | 31 | 28 |
| PFN1 | Profilin-1 | 32 | 38 |
| RAP1B | Ras-related protein Rap-1b | 33 | 48 |
| ITGB1 | Integrin beta-1 | 34 | 44 |
| HSPA5 | 78 kDa glucose-regulated protein | 35 | 83 |
| SLC3A2 | 4F2 cell-surface antigen heavy chain | 36 | 36 |
| HIST1H4A | Histone H4 | 37 | 56 |
| GNB2 | Guanine nucleotide-binding protein G(I)/G(S)/G(T) subunit beta-2 | 38 | 51 |
| ATP1A1 | Sodium/potassium-transporting ATPase subunit alpha-1 | 39 | 27 |
| YWHAQ | 14-3-3 protein theta | 40 | 40 |
| FLOT1 | Flotillin-1 | 41 | 33 |
| FLNA | Filamin-A | 42 | 67 |
| CLIC1 | Chloride intracellular channel protein 1 | 43 | 42 |
| CCT2 | T-complex protein 1 subunit beta | 44 | 49 |
| CDC42 | Cell division control protein 42 homolog | 45 | 47 |
| YWHAG | 14-3-3 protein gamma;14-3-3 protein gamma, N-terminally processed | 46 | 50 |
| GNAI2 | Guanine nucleotide-binding protein G(i) subunit alpha-2 | 52 | 41 |
| ANXA1 | Annexin A11 | 53 | 43 |
| RHOA | Transforming protein RhoA | 54 | 74 |
| PRDX2 | Peroxiredoxin-2 | 56 | 73 |
| GDI2 | Rab GDP dissociation inhibitor beta | 57 | 66 |
| ACTN4 | Alpha-actinin-4 | 59 | 52 |
| YWHAB | 14-3-3 protein beta/alpha;14-3-3 protein beta/alpha, N-terminally processed | 60 | 34 |
| RAB7A | Ras-related protein Rab-7a | 61 | 81 |
| LDHB | L-lactate dehydrogenase B chain;L-lactate dehydrogenase | 62 | 35 |
| GNAS | Guanine nucleotide-binding protein G(s) subunit alpha isoforms short | 63 | 69 |
| RAB5C | Ras-related protein Rab-5C | 64 | 53 |
| ANXA6 | Annexin A6;Annexin | 66 | 32 |
| ANXA11 | Annexin A11 | 67 | 97 |
| KPNB1 | Importin subunit beta-1 | 69 | 57 |
| EZR | Ezrin | 70 | 30 |
| ACLY | ATP-citrate synthase | 72 | 64 |
| TFRC | Transferrin receptor protein 1 | 74 | 89 |
| GNB1 | Guanine nucleotide-binding protein G(I)/G(S)/G(T) subunit beta-1 | 77 | 37 |
| RAN | GTP-binding nuclear protein Ran | 79 | 72 |
| CCT3 | T-complex protein 1 subunit gamma | 83 | 75 |
| AHCY | Adenosylhomocysteinase | 84 | 59 |
| UBA1 | Tubulin alpha-1A chain;Tubulin alpha-3C/D chain;Tubulin alpha-3E chain | 85 | 68 |
| BSG | Basigin | 91 | 61 |
| TCP1 | T-complex protein 1 subunit alpha | 95 | 78 |
| MYH9 | Myosin-9 | 99 | 29 |

**Figure S3.**
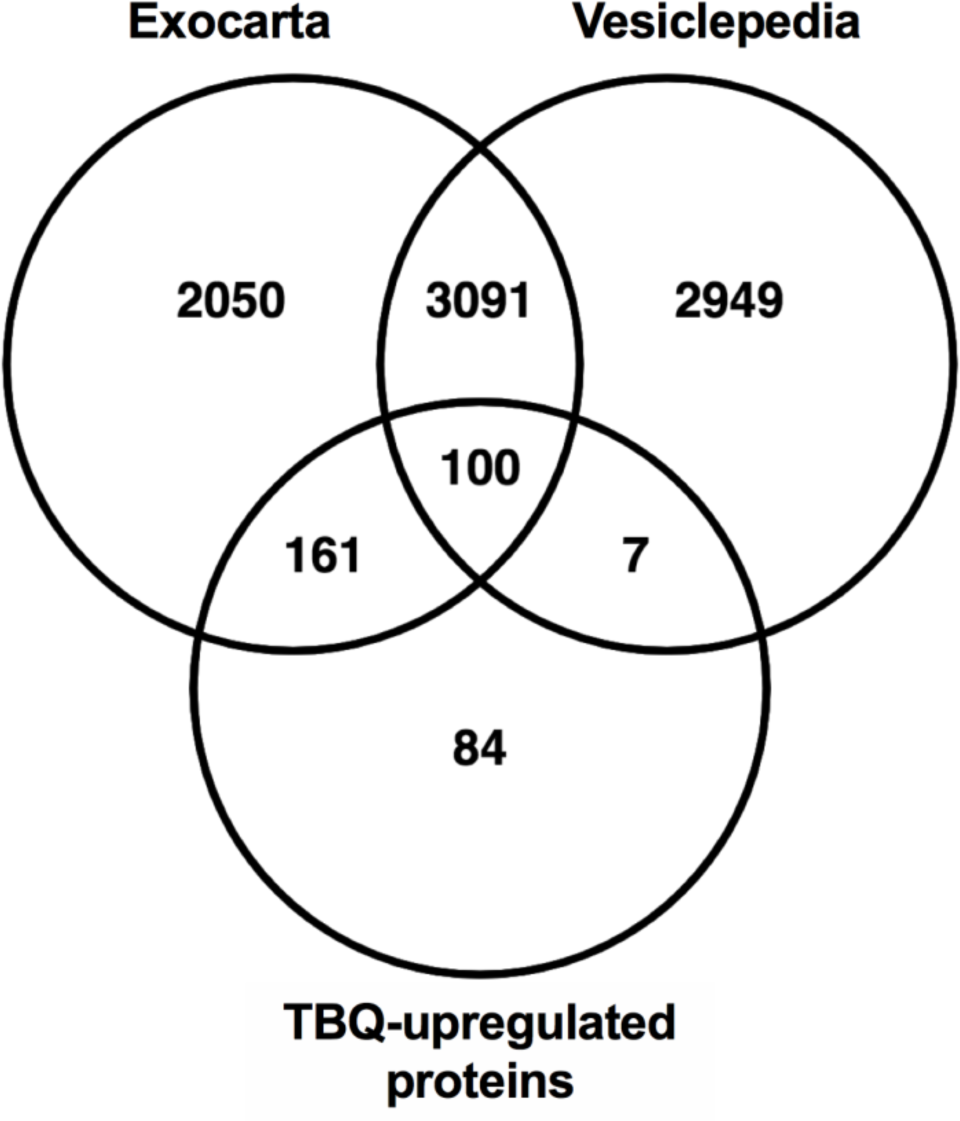
(Related to Fig. 2) TBQ-upregulated proteins share similarity with Exocarta and Vesiclepedia but contain 84 unique proteins. Venn diagram of TBQ-upregulated proteins (*e.g*., the proteins that are significantly upregulated in necroptotic vs. control EVs, with a FDR cutoff of 0.1 and S0 cutoff of 0.1) compared with the exosome proteome data bases, Exocarta and Vesiclepedia.

**Table S2.** (Related to Fig. 2) LFQ intensity of death-inducing signaling complex (DISC) components in the extracted EVs.

| Gene Name | LFQ intensity - None |  |  |  |  |  | LFQ intensity - TBQ |  |  |  |  |  | Fold change (TBQ/None) | T-test q-value |
| --- | --- | --- | --- | --- | --- | --- | --- | --- | --- | --- | --- | --- | --- | --- |
|  | sz168 | sz170 | sz171 | sz186 | sz187 | sz189 | sz168 | sz170 | sz171 | sz186 | sz187 | sz189 |  |  |
| casp8 | 20.56150818 | 20.73325539 | 21.58719063 | 18.86445045 | 20.63602829 | 19.38393021 | 23.89347076 | 25.26086998 | 23.53635025 | 25.11215019 | 24.17956924 | 25.07266998 | 18.5 | 0.02 |
| FADD | 21.84625053 | 21.67556763 | 21.65538979 | 18.76533508 | 22.13015938 | 17.55722046 | 19.66062737 | 22.19426918 | 23.00528908 | 22.47507095 | 21.55036926 | 21.44597054 | 2.16 | 0.37 |
| TRADD | 21.14176178 | 18.02750778 | 18.49320412 | 18.63318062 | 18.79969025 | 20.5822506 | 18.72833061 | 20.07887077 | 22.947855 | 19.0666008 | 19.47277069 | 20.67425919 | 1.84 | 0.45 |
| TRAF2 | 21.10859489 | 20.14731026 | 17.47186279 | 21.79729843 | 19.86410522 | 18.24882889 | 22.57074165 | 20.06526566 | 19.73435593 | 17.62561035 | 19.60129929 | 20.94464493 | 1 | 1 |
| TNFRSF10B | 21.67164612 | 19.8834877 | 17.70049095 | 18.16868973 | 20.30619049 | 20.28263092 | 19.10533905 | 19.79166985 | 19.50793076 | 20.3925705 | 20.7968502 | 20.31518936 | 1.24 | 0.66 |
| TNFRSF1B | 20.82457161 | 20.47545624 | 19.33423996 | 19.98062897 | 20.39915466 | 19.97661018 | 19.53918076 | 20.68142128 | 20.84360123 | 19.04958153 | 20.74773598 | 20.07275391 | 1.00 | 1.00 |
| TNFRSF1A | Nan | Nan | Nan | Nan | Nan | Nan | Nan | Nan | Nan | Nan | Nan | Nan |  |  |
| CFLAR (FLIP) | Nan | Nan | Nan | Nan | Nan | Nan | Nan | Nan | Nan | Nan | Nan | Nan |  |  |
| RIPK1 | Nan | Nan | Nan | Nan | Nan | Nan | Nan | Nan | Nan | Nan | Nan | Nan |  |  |
| RIPK3 | Nan | Nan | Nan | Nan | Nan | Nan | Nan | Nan | Nan | Nan | Nan | Nan |  |  |

**Table S3.** (Related to Fig. 3) TBQ-upregulated proteins assigned to the enriched GO and KEGG pathways. Proteins that are also present in the Vesiclepedia database are bolded.

| <u>ESCRT III complex</u> | <u>Regulation of type I interferon production</u> | <u>Antigen processing and presentation of exogenous peptide antigen via MHC class I</u> | <u>Phospholipid binding</u> | <u>Vesicle-mediated transport</u> |  |
| --- | --- | --- | --- | --- | --- |
| CHMP1A | DDX3X;DDX3Y | B2M | ANXA1 | ANXA11 | KIAA1033 |
| CHMP1B | FLOT1 | PSMC1 | ANXA11 | AP3B1 | LMAN1 |
| CHMP4B | IKBKB | PSMC2 | ANXA4 | ARF4 | LRSAM1 |
|  | ITCH | PSMC6 | ANXA6 | BCAP31 | MYO1F |
|  | POLR3A | PSMD12 | ANXA6 | CANX | MYO1G |
|  | POLR3D | PSMD3 | ANXA7 | CCDC22 | NSF |
|  | RPS27A;UBB;UBC | PSMD7 | ARHGAP9 | CHMP1A | PACSIN3 |
|  | TRIM56 | RPS27A;UBB;UBC | BTK | CHMP2A | PLIN3 |
|  |  | SEC61B | CHMP2A | CHMP4B | PPT1 |
|  |  |  | CPNE1 | CHMP5 | RAB5A |
|  |  |  | CPNE3 | COG2 | RDH11 |
|  |  |  | ESYT1 | COPA | RPS27A;UBB;UBC |
|  |  |  | ESYT2 | COPB1 | RTN3 |
|  |  |  | IQGAP2 | CPNE1 | SNX2 |
|  |  |  | MITD1 | CPNE3 | SNX9 |
|  |  |  | MYO1G | DENND3 | SQSTM1 |
|  |  |  | PACSIN3 | DNAJC5 | TRIM27 |
|  |  |  | SNX2 | DOCK2 | TXLNA |
|  |  |  | SNX9 | ESYT2 | VPS11 |
|  |  |  | WDFY4 | GOLGA3 | VPS4A |
|  |  |  |  | HTT | VPS4B |
|  |  |  |  | KIAA0196 | WDFY4 |

| <u>Necroptosis</u> | <u>Toll-like receptor signaling pathway</u> |
| --- | --- |
| CASP8 | BTK |
| MLKL | CASP8 |
| RPS27A;UBB;UBC | CDC2;CDK1 |
|  | IKBKB |
|  | MAP2K3 |
|  | PIK3AP1 |
|  | RPS27A;UBB;UBC |

**Table S4.**
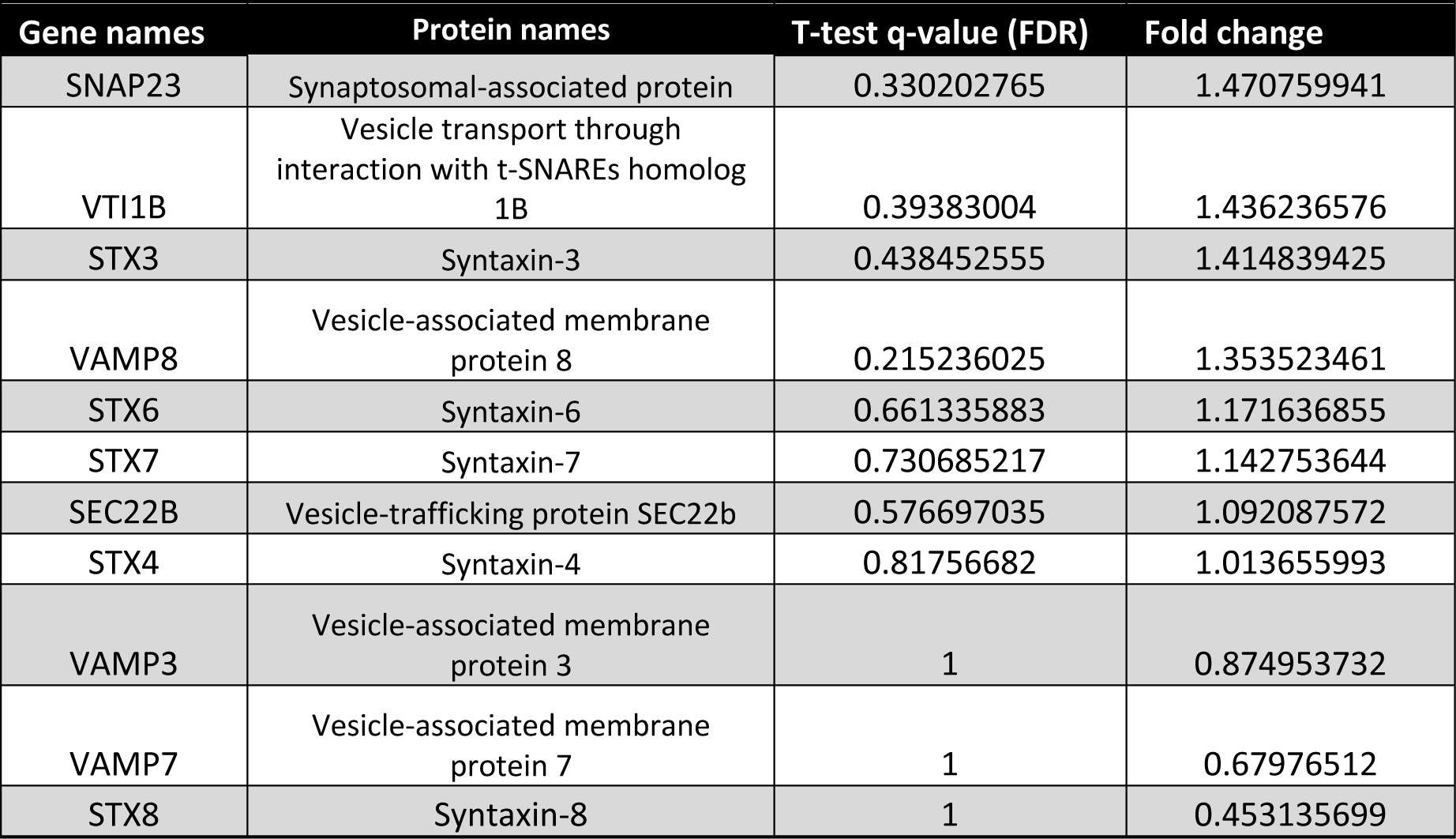
(Related to Fig. 4) SNARE proteins identified in the necroptotic EVs.

**Table S5.** (Related to Fig. 4) Rab proteins identified in increased numbers in the necroptotic EVs.

| Gene names | Protein names | T-test q-value (FDR) | Fold change |
| --- | --- | --- | --- |
| RAB5A | Ras-related protein Rab-5A | 0.087483871 | 3.70098328 |
| RAB3GAP1 | Rab3 GTPase-activating protein catalytic subunit | 0.109455471 | 2.933232655 |
| RAB3D | Ras-related protein Rab-3D | 0.197904132 | 2.445092994 |
| RABGAP1;RABGAP1L | Rab GTPase-activating protein 1;Rab GTPase-activating protein 1-like, isoform 10 | 0.074028369 | 2.277118621 |
| RAB3GAP2 | Rab3 GTPase-activating protein non-catalytic subunit | 0.152137097 | 2.276411761 |
| RAB27A | Ras-related protein Rab-27A | 0.298240786 | 2.253905732 |
| RAB6A;RAB6B | Ras-related protein Rab-6A;Ras-related protein Rab-6B | 0.334421893 | 2.02552326 |
| RAB39A | Ras-related protein Rab-39A | 0.487346251 | 2.020045913 |
| RAB4A | Ras-related protein Rab-4A | 0.301693431 | 1.750816521 |
| RABGAP1L | Rab GTPase-activating protein 1-like | 0.491986656 | 1.70643968 |
| RAB5C | Ras-related protein Rab-5C | 0.074357143 | 1.702516058 |
| RABGEF1 | Rab5 GDP/GTP exchange factor | 0.388052209 | 1.70198044 |
| RAB20 | Ras-related protein Rab-20 | 0.294641884 | 1.573590383 |
| RALA | Ras-related protein Ral-A | 0.280316883 | 1.396089652 |
| RAB11FIP1 | Rab11 family-interacting protein 1 | 0.421975309 | 1.292211209 |
| RAB18 | Ras-related protein Rab-18 | 0.663071879 | 1.264642718 |
| RAB21 | Ras-related protein Rab-21 | 0.482321155 | 1.253386823 |
| RAB27B | Ras-related protein Rab-27B | 0.544103976 | 1.227709852 |
| RAB11B | Ras-related protein Rab-11B | 0.54047619 | 1.220117196 |
| RABEP1 | Rab GTPase-binding effector protein 1 | 0.533253685 | 1.208783136 |
| RAB1B;RAB1C | Ras-related protein Rab-1B;Putative Ras-related protein Rab-1C | 0.555641566 | 1.175131154 |
| RAB9A | Ras-related protein Rab-9A | 0.615936813 | 1.126404214 |
| RAB14 | Ras-related protein Rab-14 | 0.578848092 | 1.096142107 |

## Notes

https://www.ebi.ac.uk/pride/archive/

### Competing Interest Statement

The authors have declared no competing interest.

## References

1. Vaux, D. L., Haecker, G. & Strasser, A. An Evolutionary on Apoptosis Perspective Minireview. Cell 76, 777–779 (1994).

2. Rickard, J. A. et al. RIPK1 regulates RIPK3-MLKL-driven systemic inflammation and emergency hematopoiesis. Cell 157, 1175–88 (2014).

3. Dillon, C. P. et al. RIPK1 blocks early postnatal lethality mediated by caspase- 8 and RIPK3. Cell 157, 1189–202 (2014).

4. Kaiser, W. J. et al. RIP1 suppresses innate immune necrotic as well as apoptotic cell death during mammalian parturition. Proc. Natl. Acad. Sci. 111, 7753–7758 (2014).

5. Silke, J., Rickard, J. A. & Gerlic, M. The diverse role of RIP kinases in necroptosis and inflammation. Nat. Immunol. 16, 689–697 (2015).

6. Croker, B. A. et al. Necroptosis. in Apoptosis and Beyond: The Many Ways Cells Die (ed. Radosevich, J.) 99–128 (John Wiley & Sons, Inc, 2018). doi:10.1002/9781119432463

7. Zhang, X., Dowling, J. P. & Zhang, J. RIPK1 can mediate apoptosis in addition to necroptosis during embryonic development. Cell Death Dis. 10, 1–11 (2019).

8. He, S. et al. Receptor Interacting Protein Kinase-3 Determines Cellular Necrotic Response to TNF-α. Cell 137, 1100–1111 (2009).

9. He, S., Liang, Y., Shao, F. & Wang, X. Toll-like receptors activate programmed necrosis in macrophages through a receptor-interacting kinase-3-mediated pathway. Proc. Natl. Acad. Sci. 108, 20054–20059 (2011).

10. Oberst, A. et al. Catalytic activity of the caspase-8-FLIP L complex inhibits RIPK3-dependent necrosis. Nature 471, 363–368 (2011).

11. Kaiser, W. J. et al. RIP3 mediates the embryonic lethality of caspase-8-deficient mice. Nature 471, 368–372 (2011).

12. Varfolomeev, E. E. et al. Targeted Disruption of the Mouse Caspase 8 Gene Ablates Cell Death Induction by the TNF Receptors, Fas / Apo1, and DR3 and Is Lethal Prenatally. Immunity 9, 267–276 (1998).

13. Shlomovitz, I., Zargrian, S. & Gerlic, M. Mechanisms of RIPK3-induced inflammation. Immunol. Cell Biol. 95, 166–172 (2017).

14. Newton, K. & Manning, G. Necroptosis and Inflammation. Annu. Rev. Biochem. 85, 743–763 (2016).

15. Lafont, E. et al. The linear ubiquitin chain assembly complex regulates TRAIL-induced gene activation and cell death. EMBO J. 36, 1147–1166 (2017).

16. Ting, A. T. & Bertrand, M. J. M. More to Life than NF-κB in TNFR1 Signaling. Trends Immunol. 37, 535–545 (2016).

17. Micheau, O. & Tschopp, J. Induction of TNF Receptor I-Mediated Apoptosis via Two Sequential Signaling Complexes. Cell 114, 181–190 (2003).

18. Brenner, D., Blaser, H. & Mak, T. W. Regulation of tumour necrosis factor signalling: Live or let die. Nat. Rev. Immunol. 15, 362–374 (2015).

19. Haas, T. L. et al. Recruitment of the Linear Ubiquitin Chain Assembly Complex Stabilizes the TNF-R1 Signaling Complex and Is Required for TNF-Mediated Gene Induction. Mol. Cell 36, 831–844 (2009).

20. Mahoney, D. J. et al. Both cIAP1 and cIAP2 regulate TNF -mediated NF-B activation. Proc. Natl. Acad. Sci. 105, 11778–11783 (2008).

21. Varfolomeev, E. et al. c-IAP1 and c-IAP2 are critical mediators of tumor necrosis factor α (TNFα)-induced NF-κB activation. J. Biol. Chem. 283, 24295–24299 (2008).

22. Dickens, L. S. et al. A Death Effector Domain Chain DISC Model Reveals a Crucial Role for Caspase-8 Chain Assembly in Mediating Apoptotic Cell Death. Mol. Cell 47, 291–305 (2012).

23. Majkut, J. et al. Differential affinity of FLIP and procaspase 8 for FADD’s DED binding surfaces regulates DISC assembly. Nat. Commun. 5, 1–12 (2014).

24. Schleich, K. et al. Stoichiometry of the CD95 Death-Inducing Signaling Complex: Experimental and Modeling Evidence for a Death Effector Domain Chain Model. Mol. Cell 47, 306–319 (2012).

25. Cho, Y. S. et al. Phosphorylation-Driven Assembly of the RIP1-RIP3 Complex Regulates Programmed Necrosis and Virus-Induced Inflammation. Cell 137, 1112–1123 (2009).

26. Murphy, J. M. et al. The pseudokinase MLKL mediates necroptosis via a molecular switch mechanism. Immunity 39, 443–453 (2013).

27. Sun, L. et al. Mixed lineage kinase domain-like protein mediates necrosis signaling downstream of RIP3 kinase. Cell 148, 213–227 (2012).

28. Hildebrand, J. M. et al. Activation of the pseudokinase MLKL unleashes the four-helix bundle domain to induce membrane localization and necroptotic cell death. Proc. Natl. Acad. Sci. 111, 15072–15077 (2014).

29. Dondelinger, Y. et al. MLKL Compromises Plasma Membrane Integrity by Binding to Phosphatidylinositol Phosphates. Cell Rep. 7, 971–981 (2014).

30. Cai, Z. et al. Plasma membrane translocation of trimerized MLKL protein is required for TNF-induced necroptosis. Nat. Cell Biol. 16, 55–65 (2014).

31. Shlomovitz, I. et al. Necroptosis directly induces the release of full-length biologically active IL-33 in vitro and in an inflammatory disease model. 1–16 (2018). doi:10.1111/febs.14738

32. Weinlich, R., Oberst, A., Beere, H. M. & Green, D. R. Necroptosis in development, inflammation and disease. Nat. Rev. Mol. Cell Biol. 18, 127–136 (2017).

33. Pasparakis, M. & Vandenabeele, P. Necroptosis and its role in inflammation. Nature 517, 311–320 (2015).

34. Linkermann, A. et al. Two independent pathways of regulated necrosis mediate ischemia-reperfusion injury. Proc. Natl. Acad. Sci. 110, 12024–12029 (2013).

35. Lin, J. et al. A Role of RIP3-Mediated Macrophage Necrosis in Atherosclerosis Development. Cell Rep. 3, 200–210 (2013).

36. Ouchida, A. T. et al. RIPK1 mediates axonal degeneration by promoting inflammation and necroptosis in ALS. Science (80-.). 353, 603–608 (2016).

37. Ofengeim, D. et al. Activation of Necroptosis in Multiple Sclerosis. Cell Rep. 10, 1836–1849

38. Caccamo, A. et al. Necroptosis activation in Alzheimer’s disease. 20, (2017).

39. Yuan, J., Amin, P. & Ofengeim, D. Necroptosis and RIPK1-mediated neuroinflammation in CNS diseases. Nat. Rev. Neurosci. 20, 19–33 (2018).

40. Guo, H. et al. Herpes simplex virus suppresses necroptosis in human cells. Cell Host Microbe 17, 243–251 (2015).

41. Huang, Z. et al. RIP1/RIP3 binding to HSV-1 ICP6 initiates necroptosis to restrict virus propagation in mice. Cell Host Microbe 17, 229–242 (2015).

42. Wang, X. et al. Direct activation of RIP3/MLKL-dependent necrosis by herpes simplex virus 1 (HSV-1) protein ICP6 triggers host antiviral defense. Proc Natl Acad Sci U S A 111, 15438–15443 (2014).

43. Koehler, H. et al. Inhibition of DAI-dependent necroptosis by the Z-DNA binding domain of the vaccinia virus innate immune evasion protein, E3. Proc. Natl. Acad. Sci. 114, 11506–11511 (2017).

44. Upton, J. W., Kaiser, W. J. & Mocarski, E. S. Cytomegalovirus M45 cell death suppression requires receptor-interacting protein (RIP) homotypic interaction motif (RHIM)-dependent interaction with RIP1. J. Biol. Chem. 283, 16966–16970 (2008).

45. Upton, J. W., Kaiser, W. J. & Mocarski, E. S. Virus inhibition of RIP3-dependent necrosis. Cell Host Microbe 7, 302–313 (2010).

46. Liu, X. et al. Epstein-Barr virus encoded latent membrane protein 1 suppresses necroptosis through targeting RIPK1/3 ubiquitination. Cell Death Dis. 9, 1–14 (2018).

47. Thapa, R. J. et al. DAI Senses Influenza A Virus Genomic RNA and Activates RIPK3-Dependent Cell Death. Cell Host Microbe 20, 674–681 (2016).

48. Kuriakose, T. et al. ZBP1/DAI is an innate sensor of influenza virus triggering the NLRP3 inflammasome and programmed cell death pathways. Sci. Immunol. 1, aag2045 (2016).

49. Robinson, N. et al. Type i interferon induces necroptosis in macrophages during infection with Salmonella enterica serovar Typhimurium. Nat. Immunol. 13, 954–962 (2012).

50. Roca, F. J. & Ramakrishnan, L. TNF dually mediates resistance and susceptibility to mycobacteria via mitochondrial reactive oxygen species. Cell 153, 521–534 (2013).

51. González-Juarbe, N. et al. Pore-Forming Toxins Induce Macrophage Necroptosis during Acute Bacterial Pneumonia. PLoS Pathog. 11, 1–23 (2015).

52. Kitur, K. et al. Toxin-Induced Necroptosis Is a Major Mechanism of Staphylococcus aureus Lung Damage. PLoS Pathog. 11, 1–20 (2015).

53. Ahn, D. & Prince, A. Participation of Necroptosis in the Host Response to Acute Bacterial Pneumonia. J. Innate Immun. 9, 262–270 (2017).

54. Aaes, T. L. et al. Vaccination with Necroptotic Cancer Cells Induces Efficient Anti-tumor Immunity. Cell Rep. 15, 274–287 (2016).

55. Aaes, T. L. et al. Immunodominant AH1 Antigen-Deficient Necroptotic, but Not Apoptotic, Murine Cancer Cells Induce Antitumor Protection. J. Immunol. 204, 775–787 (2020).

56. Lin, S. Y. et al. Necroptosis promotes autophagy-dependent upregulation of DAMP and results in immunosurveillance. Autophagy 14, 778–795 (2018).

57. Snyder, A. G. et al. Intratumoral activation of the necroptotic pathway components RIPK1 and RIPK3 potentiates antitumor immunity. Sci. Immunol. 4, (2019).

58. Van Lint, S. et al. Treatment with mRNA coding for the necroptosis mediator MLKL induces antitumor immunity directed against neo-epitopes. Nat. Commun. 9, 1–17 (2018).

59. Yatim, N. et al. RIPK1 and NF-kB signaling in dying cells determines cross-priming of CD8+ T cells. Science (80-.). 350, 328–334 (2015).

60. Sebbagh, M. et al. Caspase-3-mediated cleavage of ROCK I induces MLC phosphorylation and apoptotic membrane blebbing. Nat. Cell Biol. 3, 346–352 (2001).

61. Martin, S. J., Green, D. R. & Cotter, T. G. Dicing with death: dissecting the components of the apoptosis machinery. Trends Biochem. Sci. 19, 26–30 (1994).

62. Martin, S. J. & Green, D. R. Protease activation during apoptosis: Death by a thousand cuts? Cell 82, 349–352 (1995).

63. Majno, G. & Joris, I. Apoptosis, oncosis, and necrosis. An overview of cell death (1995, Am. J. Pathol.).pdf. Am. J. Pathol. 146, 3–15 (1995).

64. Zargarian, S. et al. Phosphatidylserine externalization, “necroptotic bodies” release, and phagocytosis during necroptosis. PLoS Biol. 15, 1–23 (2017).

65. Gong, Y. N. et al. ESCRT-III Acts Downstream of MLKL to Regulate Necroptotic Cell Death and Its Consequences. Cell 169, 286-300.e16 (2017).

66. Yoon, S. et al. MLKL, the Protein that Mediates Necroptosis, Also Regulates Endosomal Trafficking and Extracellular Vesicle Generation. Immunity 47, 51-65.e7 (2017).

67. Williams, R. L. & Urbé, S. The emerging shape of the ESCRT machinery. Nat. Rev. Mol. Cell Biol. 8, 355–368 (2007).

68. Gong, Y. N., Guy, C., Crawford, J. C. & Green, D. R. Biological events and molecular signaling following MLKL activation during necroptosis. Cell Cycle 16, 1748–1760 (2017).

69. Van Niel, G., D’Angelo, G. & Raposo, G. Shedding light on the cell biology of extracellular vesicles. Nat. Rev. Mol. Cell Biol. 19, 213–228 (2018).

70. Johnstone, R. M., Adam, M., Hammond, J. R., Orr, L. & Turbide, C. Vesicle formation during reticulocyte maturation. Association of plasma membrane activities with released vesicles (exosomes). J. Biol. Chem. 262, 9412–9420 (1987).

71. Pan, B. T., Teng, K., Wu, C., Adam, M. & Johnstone, R. M. Electron microscopic evidence for externalization of the transferrin receptor in vesicular form in sheep reticulocytes. J. Cell Biol. 101, 942–948 (1985).

72. Butler, W. T. The nature and significance of osteopontin. Connect. Tissue Res. 23, 123–136 (1989).

73. Tricarico, C., Clancy, J. & D’Souza-Schorey, C. Biology and biogenesis of shed microvesicles. Small GTPases 8, 220–232 (2017).

74. Zhang, H. et al. Identification of distinct nanoparticles and subsets of extracellular vesicles by asymmetric flow field-flow fractionation. Nat. Cell Biol. 20, 332–343 (2018).

75. Colombo, M., Raposo, G. & Théry, C. Biogenesis, Secretion, and Intercellular Interactions of Exosomes and Other Extracellular Vesicles. Annu. Rev. Cell Dev. Biol. 30, 255–289 (2014).

76. Sundström, C. & Nilsson, K. Establishment and characterization of a human histiocytic lymphoma cell line (U-937). Int. J. Cancer 17, 565–577 (1976).

77. Tyanova, S., Temu, T. & Cox, J. The MaxQuant computational platform for mass spectrometry-based shotgun proteomics. Nat. Protoc. 11, 2301–2319 (2016).

78. Cox, J. & Mann, M. MaxQuant enables high peptide identification rates, individualized p.p.b.-range mass accuracies and proteome-wide protein quantification. Nat. Biotechnol. 26, 1367–1372 (2008).

79. Cox, J. et al. Accurate proteome-wide label-free quantification by delayed normalization and maximal peptide ratio extraction, termed MaxLFQ. Mol. Cell. Proteomics 13, 2513–2526 (2014).

80. Tyanova, S. et al. The Perseus computational platform for comprehensive analysis of (prote)omics data. Nat. Methods 13, 731–740 (2016).

81. Oliveros, J. C. Venny. An interactive tool for comparing lists with Venn’s diagrams. Available at: https://bioinfogp.cnb.csic.es/tools/venny/index.html.

82. Mathivanan, S., Fahner, C. J., Reid, G. E. & Simpson, R. J. ExoCarta 2012: Database of exosomal proteins, RNA and lipids. Nucleic Acids Res. 40, 1241–1244 (2012).

83. Simpson, R. J., Kalra, H. & Mathivanan, S. Exocarta as a resource for exosomal research. J. Extracell. Vesicles 1, (2012).

84. Keerthikumar, S. et al. ExoCarta: A Web-Based Compendium of Exosomal Cargo. J. Mol. Biol. 428, 688–692 (2016).

85. Pathan, M. et al. Vesiclepedia 2019: A compendium of RNA, proteins, lipids and metabolites in extracellular vesicles. Nucleic Acids Res. 47, D516–D519 (2019).

86. Kalra, H. et al. Vesiclepedia: A Compendium for Extracellular Vesicles with Continuous Community Annotation. PLoS Biol. 10, 8–12 (2012).

87. Ritchie, M. E. et al. Limma powers differential expression analyses for RNA-sequencing and microarray studies. Nucleic Acids Res. 43, e47 (2015).

88. Shannon, P. et al. Cytoscape: A Software Environment for Integrated Models. Genome Res. 13, 426 (1971).

89. Deutsch, E. W. et al. The ProteomeXchange consortium in 2020: enabling ‘big data’ approaches in proteomics. Nucleic Acids Res. 48, D1145–D1152 (2020).

90. Jones, P. et al. PRIDE: New developments and new datasets. Nucleic Acids Res. 36, 878–883 (2008).

91. Perez-Riverol, Y. et al. The PRIDE database and related tools and resources in 2019: Improving support for quantification data. Nucleic Acids Res. 47, D442–D450 (2019).

92. Fadok, V. A. et al. Exposure of phosphatidylserine on the surface of apoptotic lymphocytes triggers specific recognition and removal by macrophages. J. Immunol. 148, 2207–16 (1992).

93. Shlomovitz, I., Speir, M. & Gerlic, M. Flipping the dogma - phosphatidylserine in non-apoptotic cell death. Cell Commun. Signal. 17, 139 (2019).

94. Edry-botzer, L. & Gerlic, M. Exploding the necroptotic bubble. Cell Strees 1, 107–109 (2017).

95. Lötvall, J. et al. Minimal experimental requirements for definition of extracellular vesicles and their functions: A position statement from the International Society for Extracellular Vesicles. J. Extracell. Vesicles 3, 1–6 (2014).

96. Van Deun, J. et al. EV-TRACK: Transparent reporting and centralizing knowledge in extracellular vesicle research. Nat. Methods 14, 228–232 (2017).

97. Doyle, L. M. & Wang, M. Z. Overview of Extracellular Vesicles, Their Origin, Composition, Purpose, and Methods for Exosome Isolation and Analysis. Cells 8, 1–24 (2019).

98. Yáñez-Mó, M. et al. Biological properties of extracellular vesicles and their physiological functions. J. Extracell. Vesicles 4, 1–60 (2015).

99. Lo Cicero, A., Stahl, P. D. & Raposo, G. Extracellular vesicles shuffling intercellular messages: For good or for bad. Curr. Opin. Cell Biol. 35, 69–77 (2015).

100. Fu, T. M. et al. Cryo-EM Structure of Caspase-8 Tandem DED Filament Reveals Assembly and Regulation Mechanisms of the Death-Inducing Signaling Complex. Mol. Cell 64, 236–250 (2016).

101. Stuermer, C. A. O. et al. Glycosylphosphatidyl Inositol-anchored Proteins and fyn Kinase Assemble in Noncaveolar Plasma Membrane Microdomains Defined by Reggie-1 and -2. Mol. Biol. Cell 12, 3031–3045 (2001).

102. Lang, D. M. et al. Identification of reggie-1 and reggie-2 as plasmamembrane-associated proteins which cocluster with activated GPI-anchored cell adhesion molecules in non-caveolar micropatches in neurons. J. Neurobiol. 37, 502–523 (1998).

103. Babuke, T. et al. Hetero-oligomerization of reggie-1/flotillin-2 and reggie-2/flotillin-1 is required for their endocytosis. Cell. Signal. 21, 1287–1297 (2009).

104. Simons, K. & Ikonen, E. Functional rafts in cell membranes. Nature 387, 569–572 (1997).

105. Simons, K. & Sampaio, J. L. Membrane organization and lipid rafts. Cold Spring Harb. Perspect. Biol. 3, 1–17 (2011).

106. Fan, W. et al. Flotillin-mediated endocytosis and ALIX–syntenin-1–mediated exocytosis protect the cell membrane from damage caused by necroptosis. Sci. Signal. 12, (2019).

107. Mei, K. & Guo, W. The exocyst complex. Curr. Biol. 28, R922–R925 (2018).

108. Cai, H., Reinisch, K. & Ferro-Novick, S. Coats, Tethers, Rabs, and SNAREs Work Together to Mediate the Intracellular Destination of a Transport Vesicle. Dev. Cell 12, 671–682 (2007).

109. Wu, B. & Guo, W. The exocyst at a glance. J. Cell Sci. 128, 2957–2964 (2015).

110. Guo, W., Roth, D., Walch-Solimena, C. & Novick, P. The exocyst is an effector for Sec4P, targeting secretory vesicles to sites of exocytosis. EMBO J. 18, 1071–1080 (1999).

111. Wu, S., Mehta, S. Q., Pichaud, F., Bellen, H. J. & Quiocho, F. A. Sec15 interacts with Rab11 via a novel domain and affects Rab11 localization in vivo. Nat. Struct. Mol. Biol. 12, 879–885 (2005).

112. Dirac-Svejstrup, A. B., Sumizawa, T. & Pfeffer, S. R. Identification of a GDI displacement factor that releases endosomal Rab GTPases from Rab-GDI. EMBO J. 16, 465–472 (1997).

113. Fauster, A. et al. Systematic genetic mapping of necroptosis identifies SLC39A7 as modulator of death receptor trafficking. Cell Death Differ. 26, 1138–1155 (2019).

114. Zaman, M. M.-U. et al. Ubiquitination-Deubiquitination by the TRIM27-USP7 Complex Regulates Tumor Necrosis Factor Alpha-Induced Apoptosis. Mol. Cell. Biol. 33, 4971–4984 (2013).

115. Matsusuhima Kouji, Larsen Christian G., Dubois, G. C. & Oppenheim, J. J. Purification and Characterization of a Novel Monocyte Chemotactic and Activating Factor. Cancer Res. 169, 1485–1490 (1989).

116. Yoshimura, T., Robinson, E. A., Tanaka, S., Appella, E. & Leonard, E. J. Purification and amino acid analysis of two human monocyte chemoattractants produced by phytohemagglutinin-stimulated human blood mononuclear leukocytes. J. Immunol. 142, 1956–62 (1989).

117. Yoshimura, T. et al. Purification and amino acid analysis of two human glioma-derived monocyte chemoattractants. J. Exp. Med. 169, 1449–1459 (1989).

118. Kreuzaler, P. & Watson, C. J. Killing a cancer: What are the alternatives? Nature Reviews Cancer (2012). doi:10.1038/nrc3264

119. Su, Z., Yang, Z., Xie, L., Dewitt, J. P. & Chen, Y. Cancer therapy in the necroptosis era. 1–9 (2016). doi:10.1038/cdd.2016.8

120. Morelli, A. E. et al. Endocytosis, intracellular sorting, and processing of exosomes by dendritic cells. Blood 104, 3257–3266 (2004).

121. Mallegol, J. et al. T84-Intestinal Epithelial Exosomes Bear MHC Class II/Peptide Complexes Potentiating Antigen Presentation by Dendritic Cells. Gastroenterology 132, 1866–1876 (2007).

122. Kreiter, S. et al. Mutant MHC class II epitopes drive therapeutic immune responses to cancer. Nature 520, 692–696 (2015).

123. Bhatnagar, S., Shinagawa, K., Castellino, F. J. & Schorey, J. S. Exosomes released from macrophages infected with intracellular pathogens stimulate a proinflammatory response in vitro and in vivo. Blood 110, 3234–3244 (2007).

124. Skokos, D. et al. Mast Cell-Derived Exosomes Induce Phenotypic and Functional Maturation of Dendritic Cells and Elicit Specific Immune Responses In Vivo. J. Immunol. 170, 3037–3045 (2003).

125. Robbins, P. D. & Morelli, A. E. Regulation of immune responses by extracellular vesicles. Nat. Rev. Immunol. 14, 195–208 (2014).

126. Holler, N. et al. Fas triggers an alternative, caspase-8-independent cell death pathway using the kinase RIP as effector molecule. Nat. Immunol. 1, 489–495 (2000).

127. Oberst, A. et al. Inducible dimerization and inducible cleavage reveal a requirement for both processes in caspase-8 activation. J. Biol. Chem. 285, 16632–16642 (2010).

128. Chinnaiyan, A. M., O’Rourke, K., Tewari, M. & Dixit, V. M. FADD, a Novel Death Domain-Containing Interacts with the Death Domain of Fas and Initiates Apoptosis. Cell 81, 505–512 (1995).

129. Kischkel, F. C. et al. Apo2L/TRAIL-dependent recruitment of endogenous FADD and caspase-8 to death receptors 4 and 5. Immunity 12, 611–620 (2000).

130. Fu, Q. et al. Structural Basis and Functional Role of Intramembrane Trimerization of the Fas/CD95 Death Receptor. Mol. Cell 61, 602–613 (2016).

131. Schleich, K. et al. Molecular architecture of the DED chains at the DISC: Regulation of procaspase-8 activation by short DED proteins c-FLIP and procaspase-8 prodomain. Cell Death Differ. 23, 681–694 (2016).

132. Henry, C. M. & Martin, S. J. Caspase-8 Acts in a Non-enzymatic Role as a Scaffold for Assembly of a Pro-inflammatory “FADDosome” Complex upon TRAIL Stimulation. Mol. Cell 65, 715-729.e5 (2017).

133. Medema, J. P. et al. FLICE is activated by association with the CD95 death-inducing signaling complex (DISC). EMBO J. 16, 2794–2804 (1997).

134. Hoffmann, J. C., Pappa, A., Krammer, P. H. & Lavrik, I. N. A New C-Terminal Cleavage Product of Procaspase-8, p30, Defines an Alternative Pathway of Procaspase-8 Activation. Mol. Cell. Biol. 29, 4431–4440 (2009).

135. Krueger, A., Schmitz, I., Baumann, S., Krammer, P. H. & Kirchhoff, S. Cellular FLICE-inhibitory Protein Splice Variants Inhibit Different Steps of Caspase-8 Activation at the CD95 Death-inducing Signaling Complex. J. Biol. Chem. 276, 20633–20640 (2001).

136. Tanzer, M. C. et al. Quantitative and Dynamic Catalogs of Proteins Released during Apoptotic and Necroptotic Cell Death. Cell Rep. 30, 1260-1270.e5 (2020).

137. Cai, Z. et al. Activation of cell-surface proteases promotes necroptosis, inflammation and cell migration. Cell Res. 26, 886–900 (2016).

138. Creutz, C. E. The annexins and exocytosis. Science (80-.). 258, 924–931 (1992).

139. Théry, C. et al. Minimal information for studies of extracellular vesicles 2018 (MISEV2018): a position statement of the International Society for Extracellular Vesicles and update of the MISEV2014 guidelines. J. Extracell. Vesicles 7, (2018).

140. Barrès, C. et al. Galectin-5 is bound onto the surface of rat reticulocyte exosomes and modulates vesicle uptake by macrophages. Blood 115, 696–705 (2010).

141. Wei, H., Malcor, J. D. M. & Harper, M. T. Lipid rafts are essential for release of phosphatidylserine-exposing extracellular vesicles from platelets. Sci. Rep. 8, 1–11 (2018).

142. Salzer, U., Hinterdorfer, P., Hunger, U., Borken, C. & Prohaska, R. Ca++-dependent vesicle release from erythrocytes involves stomatin-specific lipid rafts, synexin (annexin VII), and sorcin. Blood 99, 2569–2577 (2002).

143. Burger, D. et al. Endothelial microparticle formation by angiotensin II is mediated via ang II receptor type I/NADPH Oxidase/rho kinase pathways targeted to lipid rafts. Arterioscler. Thromb. Vasc. Biol. 31, 1898–1907 (2011).

144. Del Conde, I., Shrimpton, C. N., Thiagarajan, P. & López, J. A. Tissue-factor-bearing microvesicles arise from lipid rafts and fuse with activated platelets to initiate coagulation. Blood 106, 1604–1611 (2005).

145. Segawa, K. & Nagata, S. An Apoptotic ‘Eat Me’ Signal: Phosphatidylserine Exposure. Trends Cell Biol. 25, 639–650 (2015).

146. Nagata, S. Apoptosis and Clearance of Apoptotic Cells. Annu. Rev. Immunol. 36, 489–517 (2018).

147. Wu, N. et al. Critical Role of Lipid Scramblase TMEM16F in Phosphatidylserine Exposure and Repair of Plasma Membrane after Pore Formation. Cell Rep. 30, 1129-1140.e5 (2020).

148. Wei, Y. et al. Pyruvate kinase type M2 promotes tumour cell exosome release via phosphorylating synaptosome-associated protein 23. Nat. Commun. 8, 1–12 (2017).

149. Jahn, R. & Scheller, R. H. SNAREs - Engines for membrane fusion. Nat. Rev. Mol. Cell Biol. 7, 631–643 (2006).

150. Fader, C. M., Sánchez, D. G., Mestre, M. B. & Colombo, M. I. TI-VAMP/VAMP7 and VAMP3/cellubrevin: two v-SNARE proteins involved in specific steps of the autophagy/multivesicular body pathways. Biochim. Biophys. Acta - Mol. Cell Res. 1793, 1901–1916 (2009).

151. Zhu, K. et al. Necroptosis promotes cell-autonomous activation of proinflammatory cytokine gene expression article. Cell Death Dis. 9, (2018).

152. Kearney, C. J. et al. Necroptosis suppresses inflammation via termination of TNF-or LPS-induced cytokine and chemokine production. Cell Death Differ. 22, 1313–1327 (2015).

